# Heterogeneous and conserved radiation responses reveal FOXM1-dependent regulation of microcephaly genes in glioblastoma

**DOI:** 10.64898/2026.09.01.748488

**Authors:** L. Van Eupen, E. Etlioglu, K. Tabury, K.L. Ligon, I. Aguirre Alcolea, M. Derweduwe, A. Claeys, A. Puspitasari-Kokko, I. Primac, F. De Smet, R. Quintens

**Affiliations:** Radiobiology Unit, Nuclear Medical Applications Institute, SCK CEN, Mol, Belgium; Laboratory for Precision Cancer Medicine, Translational Cell and Tissue Research Unit, Department of Imaging and Pathology, KU Leuven, Leuven, Belgium; Leuven Institute for Single-Cell Omics (LISCO), KU Leuven, 3000 Leuven, Belgium; Leuven Cancer Institute (LKI), KU Leuven, 3000 Leuven, Belgium; Center for Neuro-oncology, Dana-Farber Cancer Institute, Boston, MA, USA; Department of Pathology, Brigham and Women’s Hospital, Boston, MA, USA; Department of Pathology, Harvard Medical School, Boston, MA, USA; GSI Helmholtzzentrum für Schwerionenforschung, Biophysics Department, Darmstadt, Germany

**Keywords:** Glioblastoma, Photon irradiation, Carbon ion irradiation, Radiation response, Microcephaly, FOXM1

## Abstract

Glioblastoma (GBM) is characterized by marked heterogeneity, glioma stem-like cells (GSCs), and resistance to therapy. Because GSCs share features with neural progenitor cells (NPCs), we investigated whether neurodevelopmental programs contribute to their response to irradiation. Transcriptional profiling of four patient-derived GSC lines revealed cell line-specific responses, with radiosensitivity correlating with the magnitude of p53 activation and basal expression of its negative regulator, MDM2. Despite this heterogeneity, radiation consistently activated p53-dependent pathways and suppressed cell-cycle programs. Among these, genes associated with primary hereditary microcephaly (MCPH) that regulate NPC proliferation were coordinately repressed. Single-cell RNA sequencing localized this response to G2/M-cycling cells. FOXM1 was similarly reduced following irradiation, emerged as a candidate regulator of a subset of MCPH genes, and correlated with their expression in GBM tumors. Pharmacological inhibition of FOXM1 reduced expression of selected MCPH genes and enhanced radiosensitivity in U251 cells. Together, these findings identify coordinated suppression of a FOXM1-associated MCPH program as part of the GBM radiation response, while suggesting that the radiosensitizing effects of pharmacological FOXM1inhibition extend beyond this transcriptional axis.

## Introduction

With a median survival of less than two years, glioblastoma (GBM, grade IV astrocytoma IDH_wt_)[1] is one of the most aggressive and lethal cancers in adults. The standard treatment regimen consists of maximal safe surgical resection, radiotherapy using photon irradiation (PIR) and concomitant temozolomide, followed by adjuvant temozolomide – a protocol that has not changed in over 20 years [2], and only prolongs survival in the range of months with nearly all patients showing relapse.

A key hallmark of GBM tumors is their very high inter- and intra-tumor heterogeneity [3]. Firstly, bulk genomic and transcriptomic analyses identified inter-patient heterogeneity leading to the current TCGA classification, including proneural (PN), classical (CL) or mesenchymal (MES) GBM [4], based on the work of Verhaak *et al.* (2010) [5]. However, single-cell profiling revealed GBM tumors as mixtures of these subtypes (intratumoral heterogeneity) [6]. Moreover, recent studies reported that GBM cells can exist in different dynamic cellular states, including axes based on developmental-like states [7], stemness [8, 9] or metabolism [8], among others. Arguably, the most commonly adopted classification frameworks in the field of GBM come from a study that identified states reflecting normal neuronal development, including oligodendrocyte precursor cell-like (OPC-like), neuronal progenitor cell-like (NPC-like), astrocyte-like (AC-like), and mesenchymal-like (MES-like) cells [10]. These signatures have recently been further refined and supplied with additional signatures comprising an even broader spectrum of phenotypic states including signatures reflecting states of stress, hypoxia and cycling cells [11].

As radiotherapy is a cornerstone of GBM treatment, the response of GBM cells to PIR has been extensively studied. In general, PIR induces DNA damage, particularly double-strand breaks, which activate the DNA damage response through ATM/ATR signaling and downstream effectors such as CHK1/CHK2 and p53. This leads to apoptosis, cell-cycle arrest and activation of DNA repair pathways, including non-homologous end joining and homologous recombination, enabling tumor cell survival and recovery [12]. A major contributor to intrinsic radioresistance is the presence of glioma stem cells (GSCs), which exhibit enhanced DNA repair capacity [12] and selective activation of DDR pathways [13], allowing them to survive treatment and repopulate the tumor [14]. For example, GSCs are significantly enriched for RAD51, which has a central role in homologous recombination, and inhibition of RAD51 results in increased radiosensitivity [15, 16]. Beyond DNA repair, GSCs adapt to radiation through dynamic changes in cell-cycle regulation [17, 18], and the tumor microenvironment [19, 20]. The plastic abilities of GSCs, and to a lesser extent those of less stem cell-like GBM cells, further support radioresistance, with distinct cellular states responding differently to radiation [21]. Mesenchymal-like phenotypes are often considered the most radioresistant due to enhanced DNA repair, hypoxia adaptation, and pro-survival signaling, whereas proneural states are typically more radiosensitive [6, 22]. Radiation can induce both selection of the resistant cells and shifting of sensitive cells to resistant phenotypes [23]. However, there is also growing evidence that metabolic and hypoxia-adapted phenotypes, cutting across subtype boundaries, can dominate radio resistance, suggesting that functional states (*e.g.,* hypoxic, immune-evasive) may be as important as classical subtype classifications [21, 24].

Unfortunately, this radioresistance of GSCs is reflected by the dismal prognosis for GBM patients indicating the current standard-of-care is a suboptimal treatment strategy, especially when considering the vast heterogeneity in the patient population. Therefore, the need for more in-depth investigations into the effects of radiation on GSCs and the development of novel treatment strategies remains imperative. In addition to PIR, other external beam radiation modalities exist for cancer treatment, including carbon ion radiation (CIR). In contrast to PIR, CIR is a high-linear energy transfer (LET) radiation that induces more complex double-stranded DNA breaks, resulting in greater biological effectiveness. Preclinical studies have shown that this leads to more GSC cell death, reduced GBM cell migration, and potentially lowered radioresistance in GSCs [25, 26]. However, more research is needed on the molecular response of heterogeneous GSCs to high-LET radiation, while infrastructural challenges and suboptimal clinical trial data limit its clinical application [27].

Considering the GSC’s radiation response, it is also important to reflect on the origin and progression of GBM. Glioma genesis parallels many aspects of normal brain development, including elements of its hierarchical structure [28, 29]. However, GBM tumor cells exhibit far greater cellular diversity and plasticity. Unlike neural stem cells, which irreversibly differentiate into cells such as neurons and glial cells, GBM cells retain the ability to revert to a more stem-like condition even after adopting differentiated-like states. Moreover, this phenotypic shifting can occur in response to stressors such as radiation [23]. Strikingly, isolation and separate culturing of different GBM cell subpopulations highlighted their individual tumor-initiating potential when each subpopulation was able to induce new tumors resembling the original tumors’ heterogeneity [10].

The similarity with normal brain development has sparked new ideas for treatment strategies [30]. For example, by targeting neurodevelopmental signaling pathways that sustain radial glia proliferation and self-renewal such as Notch, WNT, SHH, BMP and FGF/ERK, which have shown to be effective in preclinical studies [31, 32]. Alternatively, targeting neurodevelopmental migration by inhibiting RhoA-ROCK or CDK5 *in vitro* and *in vivo* has been shown to diminish GBM invasion [28]. Another extensively studied treatment strategy, not only for GBM, but also for many different cancers, is targeting the cell cycle [33]. Dysregulated cell-cycle machinery is considered to be a universal driver of cancer proliferation and targeting its components has shown promising results, even in a clinical setting . For example, inhibitors of cyclin-dependent kinases are used to treat hormone receptor-positive, HER2-negative advanced breast cancer [34] and have been considered for GBM treatment as well. Abemaciclib and AT7519 effectively induced cell death, cell cycle arrest, and tumor-suppressing micro-RNA in preclinical GBM models [35, 36], although clinical success has been rather limited [37]. Another way of targeting the cell cycle is to target the microtubules, critical components of spindle formation and chromosome segregation during mitosis, using microtubule-targeting agents (MTAs). These have been tested for central nervous system tumors in clinical trials and show promising results [38–40]. However, MTAs are not cancer cell-specific, leading to serious adverse effects, including peripheral and autonomic neuropathies and myelosuppression [41].

A more recently proposed angle for GBM treatment is targeting microcephaly primary hereditary (MCPH) genes [42]. So far, 30 MCPH genes have been identified. They play pivotal roles in DNA damage response, centrosome biogenesis, and proliferation of neural progenitor cells [43]. Mutations of these genes occur in this genetic form of microcephaly (*i.e,* abnormal brain development leading to significantly smaller head size) [43]. Interestingly, while expressed in all proliferating cells regardless of the cell type, they are only critical in neural progenitors [43]. Given the resemblance between GSCs and neural progenitors and the observed overexpression of some MCPH genes in low- and especially high-grade glioma [44], targeting these genes instead of the microtubules themselves might overcome the observed toxicity of MTAs, a strategy that has gained attention in the past decade [42]. Radiation has been observed to repress some MCPH genes *in vitro* in human neural progenitor cells via p53-E2F4/DREAM [45, 46]. MCPH genes as potential targets for GBM have been studied separately, but not as a collective targetable signature. Moreover, whether radiation similarly regulates this MCPH gene in GSCs and the regulatory mechanisms through which they respond are currently unknown.

In this study, we profiled the radiation response of heterogeneous patient-derived glioma stem-like cell lines (PDGSCLs) to PIR and CIR, ultimately exploring the potential regulatory mechanisms of radiation-induced repression of MCPH genes in GBM. Based on our findings, we propose Forkhead Box M1 (FOXM1) as one of the key players in this process. FOXM1 is a transcription factor with crucial activity in regulating the cell cycle, proliferation and DNA damage response [47] and its expression is increased in GBM, linking it to epithelial-to-mesenchymal transition (EMT), invasion, radioresistance and ultimately poor prognosis [48–50]. Moreover, preclinical studies have shown that inhibition of FOXM1, both *in vitro* and *in vivo* has resulted in increased sensitivity to chemo- and radiotherapy, decreased tumorigenicity, and prolonged survival of orthotopic xenograft mice models [49, 51]. In glioma cells, FOXM1 is known to interact with STAT proteins, Wnt/ß-Catenin, MELK, RAD51 and growth factors [47–49, 52, 53]. It also regulates transcription of ASPM, one of the MCPH genes, by binding to its promoter region in GBM and overexpression of FOXM1 favors malignant glioma progression through ASPM [54]. No such direct link between FOXM1 and other MCPH genes has been reported so far, although their involvement in cell cycle processes suggests similar regulatory relationships may exist.

## Materials and Methods

### Patient-derived cell lines

PDGSCLs derived from high-grade brain tumors were used and developed in the framework of studies S59804 and S61081 which were approved by the Ethische Commissie Onderzoek UZ/KU Leuven. Fresh high-grade brain tumor biopsies were collected from several hospitals, *e.g*., UZ Leuven (Leuven, Belgium; IRB protocol S59804), Jessa Ziekenhuis (Hasselt, Belgium; IRB B243201941451), AZNikolaas (Sint Niklaas, Belgium; IRB protocol EC18021) and PDGSCLs were generated by the LPCM (IRB protocol S59804, LBT- and CME-numbers). These cell lines were generated through culturing of dissociated patient biopsies taken during initial or recurrent tumor resection (**Figure 1A**). When cells persisted after three passages *in vitro*, they were further expanded and cryopreserved. BT360 cells originated from the Dana-Farber Cancer Institute (BT-number, Boston, MA, USA) in 2017 (IRB 10–417). All four patients were diagnosed with glioblastoma multiforme (GBM, WHO classification 2021); other patient demographics are shown in **Table S12**. Cells were cultured using serum-free medium, including DMEM-F12 + Glutamine (Life Technologies; #11320074, Thermo Fisher Scientific, Waltham, MA, USA), heparine (Stem Cell Technologies; #07980, Vancouver, Canada), Human recombinant EGF (Stem Cell Technologies; #78006.2, Vancouver, Canada), Human recombinant BFGF (Stem Cell Technologies; #78003.2, Vancouver, Canada), Antibiotic-Antimycotic 100X (Life technologies; #15240-062, Thermo Fisher Scientific, Waltham, MA, USA), and in-house-made supplements optimized for the PDGSCLs. Adherent lines (CME and LBT) were cultured using laminin coating (L2020-1MG, Merck/Sigma-Aldrich, Darmstadt, Germany), suspension lines (BT) were seeded without such coating in ultra-low attachment vessels. U-251 MG (U251) cells were obtained from Cytion (RRID: CVCL_0021, #300385, Heidelberg, Germany) and cultured in DMEM (Cytion, #820300a, Heidelberg, Germany) supplemented with 10% fetal bovine serum (GIBCO, #10270106, Thermo Fisher Scientific, Waltham, MA, USA) and 1% penicillin–streptomycin (GIBCO, #15140-122, Thermo Fisher Scientific, Waltham, MA, USA).

**Figure 1:**
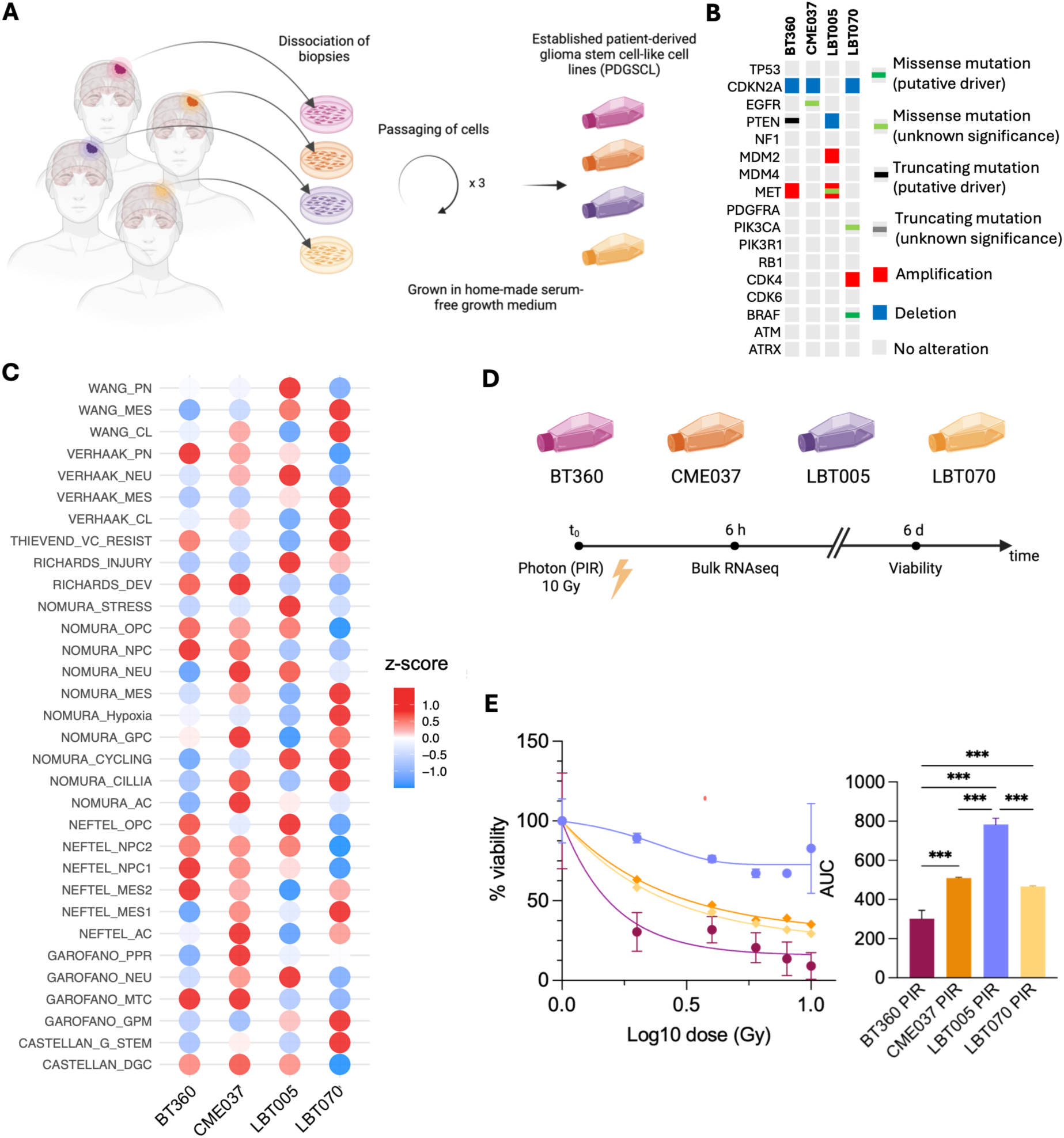
Overall sensitivity is mainly driven by cell line heterogeneity. **A**: PDGSCL generation. **B**: Whole genome sequencing data for the most common mutations, obtained from cbioportal. **C**: Enrichment analysis of four different PDGSCLs for 27 GBM phenotype signatures. Enrichment scores were calculated using ssGSEA and normalized per row to obtain z-scores. **D**: Experimental setup and timing. **E**: Dose-response curves for PIR (left) and area-under-curve values of viability measurements (right). Technical replicates = 4; error bars = SD; Ordinary one-way ANOVA with Tukey’s multiple comparisons test (alpha = 0.05); * = ≤ 0.05; ** = ≤ 0.01; *** = ≤ 0.001; **** = ≤ 0.0001.

### Irradiations

Photon irradiations (PIR; X-rays) were performed at the Stem Cell Institute Leuven (SCIL) using the RS 2000 small-animal irradiator (Rad Source, Atlanta, GA, USA) or at SCK CEN (Laboratory of Nuclear Calibration, an ISO 17025:2017 accredited facility) using an H-250 X-ray irradiator. Carbon ion irradiations (CIR) were performed at the Cave A branch of the SIS18 synchrotron at the GSI Helmholtzzentrum für Schwerionenforschung (Darmstadt, Germany) in the frame of the FAIR Phase-0 call (BioPAC 2020 project SBio08_Quintens). A 240 MeV/n ^12^C-ion beam was extracted in air and delivered using a spread-out Bragg peak (SOBP) configuration. Sham-irradiated samples served as controls in all experiments. Standard radiation doses for both PIR and CIR were 10 Gy. For viability assays, cells were irradiated with 2, 4, 6, 8 and 10 Gy.

### FOXM1 inhibition

FDI-6 binds to the FOXM1 protein, thereby inhibiting DNA binding and subsequent transcriptional regulation of its targets. FDI-6 was obtained from MedChemExpress (#HY-112721, Monmouth Junction, NJ, USA). Treatment concentrations were 20 µM, 40 µM, and 60 µM, and cells were lysed 2 and 6 hours after treatment. For combined FDI-6 and PIR experiments, increasing doses of FDI-6 (1.25, 2.5, 5, 10, 20 and 40 µM) were administered 6 hours prior to 10 Gy PIR, and cells were processed 3 days after PIR for viability measurements. Control cells were always treated with DMSO at a concentration equal to the highest FDI-6 concentration.

### Viability assays

Cells were seeded in 96-well plates (10,000 cells/well, n = 4 or n = 3). Cell viability was estimated using the ATP-based CellTiter-Glo® (Promega, Madison, WI, USA) viability assay according to the manufacturer’s instructions at six days after irradiation. Luminescence was measured using the SpectraMax® iD3 microplate reader (Molecular Devices, LLC, USA).

### Cell proliferation assay

Cells were seeded in 96-well plates (2,500 cells/well, quadruplicate). Cell proliferation was monitored using live-cell imaging with IncuCyte® Zoom 2018A Live-Cell Analysis System.

### RT-qPCR

Cells were seeded in T12.5 culture flasks (350,000 cells/flask, n = 3) or 6-well plates (800,000 cells/well, triplicate). Cells were lysed at 6 hours after irradiation, 2 and 6 hours after FDI-6 treatment and further processed using the RNeasy® Micro Plus RNA extraction kit (QIAGEN, Hilden, Germany) according to the manufacturer’s instructions. RNA concentration and purity were measured using the NanoDrop™ 2000 Spectrophotometer (Thermo Fisher Scientific, Waltham, MA, USA). cDNA synthesis was subsequently performed using the GoScript^TM^ Reverse Transcription Mix (Promega, Madison, USA). RT-qPCR was done using QIAGEN’s 2x Quantinova SYBR Green® PCR Master Mix and run on the qTOWER^3^G (Analytik Jena, Jena, Germany). *GAPDH* and *RPL13A* were used as reference genes. Primer sequences are listed in Supplementary Table S1.

### Bulk RNA sequencing

For bulk RNA sequencing, cells were seeded in T12.5 culture flasks (300,000 cells/flask, triplicate) or 6-well plates (800,000 cells/well, triplicate). RNA samples (30 ng/µl, n = 3 for PIR, n = 2 for CIR) were submitted to Novogene (Cambridge, UK) for library preparation and sequencing. Messenger RNA was purified from total RNA using poly-T oligo-attached magnetic beads. After fragmentation, the first strand cDNA was synthesized using random hexamer primers followed by the second strand cDNA synthesis. The library was ready after end repair, A-tailing, adapter ligation, size selection, amplification, and purification. The library was checked with Qubit and real-time PCR for quantification and bioanalyzer for size distribution detection. Libraries were prepared using poly(A) enrichment . Quantified libraries were pooled and enrichment and sequenced on an Illumina NovaSeq 6000 platform (Illumina, San Diego, CA, USA) as paired-end reads (150 bp), generating ∼30 million reads per sample.

### Single-cell RNA sequencing

Cells were seeded in T25 flasks (2,000,000 cells/flask) and harvested 6 hours after irradiation. Harvesting was done according to the 10x Genomics (10x Genomics, Pleasanton, CA, USA) Chromium Fixed RNA Profiling protocol (CG000478). Samples were stored at -80 °C before library prep and sequencing according to the 10x Genomics Chromium Fixed RNA Profiling Workflow for multiplexed samples (CG000527).

### Data analysis

Statistical analysis was performed using GraphPad Prism v10.0.3 (GraphPad Software, San Diego, CA, USA).

Bulk RNAseq data were processed using the nf-core rnasseq pipeline [55] and further analyzed using R (v4.3.3) and the relevant packages, such as DESeq2 (v1.42.1) [56]. Differentially expressed genes (DEGs) were computed with DESeq2 using the Wald statistic with Benjamini–Hochberg (BH) False Discovery Rate (FDR) correction. For DEGs, adjusted p-value threshold was set at 0.05; log2FoldChange (LFC) threshold was set at 0.

Rank-rank hypergeometric overlap analysis (RRHO) was performed using the RedRibbon package (v1.4) [57]. Genes were ranked according to a signed significance metric calculated as the sign of the log2 fold change multiplied by the negative log10-transformed p-value. Positive scores indicate upregulated genes, whereas negative scores indicate downregulated genes. The magnitude of the score reflects the statistical significance of differential expression.

Gene Set Enrichment Analysis (GSEA) was performed using fGSEA [58]. Genes were ranked based on LFC, and enrichment scores (ES) were calculated using the running sum (Kolmogorov-Smirnov-like) statistic, normalized for the gene set sizes (NES), and corrected for multiple testing using false discovery rate (FDR) correction (Benjamini-Hochberg; BH). Other packages for enrichment analyses included clusterProfiler (v4.10.1) [59] and enrichR (v3.2) [60]. The significance threshold for the adjusted p-value was set at 0.25.

Transcription factor (TF) activity was estimated using DoRothEA (v1.14.1) [61] and VIPER (V1.36.0) [62], and TF binding enrichment was estimated using the “ENCODE_and_ChEA_Consensus_TFs_from_ChIP-X” database from enrichR.

Average ChIP-seq signal scoring is performed on data obtained from ChIP-Atlas [63] using pyBigWig (v0.3). Gene annotations (GTF) were retrieved from GENCODE v49 (GRCh38). For each signature, the transcription start site (TSS) of each gene was identified from the GTF, and a ±2,000 bp window centered on the TSS was extracted. Mean ChIP-seq signal within each TSS window was computed. For visualization, each TSS window was divided into 100 equal bins and the mean signal per bin was retrieved; genes on the minus strand were reverse complemented so that all profiles are oriented 5′→3′. Shades correspond to mean ± SEM binned profile for each signature.

Single-cell RNAseq (scRNAseq) data was processed using the 10X Genomics CellRanger pipeline (v7.1.0) [64] and further analyzed using R (v4.3.3) and the relevant packages, such as Seurat (v5.1.0) [65]. Cells were randomly downsampled to retain 2,000 cells per condition. Cell cycle phase scores were calculated for each cell using canonical S-phase and G2/M marker genes implemented in Seurat’s CellCycleScoring() function. Cell cycle effects were mitigated by regressing S-phase and G2/M scores during normalization with SCTransform prior to downstream analyses, thereby facilitating the identification of PIR-induced transcriptional changes and biologically relevant cellular heterogeneity. Module scores for different cell states (e.g. Nomura states) and for DREAM-dependent and -independent signatures were calculated using the AddModuleScore function. Cell states were assigned based on the highest module score or classified as “mixed” when the difference between the first and second highest module score was less than 0.01. Virtual FOXM1 knockout was performed using the scTeniFoldKnk package [63].

### Public datasets

Data from the publicly available databases - The Cancer Genome Atlas (TCGA) [66] and the Chinese Glioma Genome Atlas (CGGA) [67] from GlioVis (https://gliovis.bioinfo.cnio.es) [68], the Gene Expression Profiling Interactive Analysis (GEPIA; http://gepia.cancer-pku.cn) [69], Evo-devo Mammalian Organs (https://apps.kaessmannlab.org/evodevoapp/) [70] and ChIP-Atlas (https://chip-atlas.org) [71] – was used to further structure our findings and place them in a broader context. Data were downloaded from the aforementioned websites to further incorporate into our analysis.

## Results

### Overall sensitivity, transcriptional profile and radiation response are mainly driven by cell line heterogeneity

We aimed to capture part of the heterogeneity across the GBM patient population by using four different PDGSCLs with varying genetic profiles (**Figure 1B**). GBM phenotype enrichment analysis using bulk RNAseq data suggested a heterogeneous composition of the four PDGSCLs (**Figure 1C**). Normalized enrichment scores were calculated for 27 different, but not mutually exclusive, GBM phenotype signatures from seven studies [4, 5, 7–11, 72]. Overall, compared to the other three cell lines, BT360 was negatively enriched for MES-like, stem cell-like and neuronal-like signatures. CME037 showed a similar pattern but with an enrichment for proliferative/progenitor-like genes. LBT005, on the other hand, showed enrichment of neuronal-like signatures, while MES- and CL-like signatures were mostly negatively enriched. Finally, LBT070 clearly showed enrichment for MES-, CL- and stem cell-like signatures, while (pro)neural-like signatures were negatively enriched. Together, this analysis suggested that the cell lines used represented different spectra of GBM subtypes.

Sensitivity to PIR was measured using an ATP-based viability assay 6 days after cells were irradiated with 2, 4, 6, 8 or 10 Gy (**Figure 1D**). Radiation sensitivity was different between the four cell lines, with LBT005 being most resistant and BT360 most sensitive. LBT070 and CME037 both showed intermediate sensitivity to acute PIR (**Figure 1E**).

To assess the early transcriptomic response of the four PDGSCLs to PIR, cells were irradiated with 10 Gy and harvested for bulk RNA sequencing after 6 hours. Bulk transcriptomic profiling of control and irradiated samples indicated that the overall transcriptional response was mainly driven by cell line, or patient, heterogeneity (**Figure 2A-B**). Analysis of differentially expressed genes (DEGs) for each PDGSCL comparing control samples with irradiated samples indicated that the transcriptional effect of PIR was more pronounced for BT360, LBT005 and LBT070 (1957, 881 and 2308 significant DEGs, respectively) compared to CME037 (135 DEGs), while the magnitude of effect was considerably smaller in LBT005 (**Figure 2C**). Enrichment analysis based on these DEGs confirmed concerted upregulation of the hallmark p53 pathway and downregulation of proliferation (**Figure 2D & Table S2**). Hallmark radiation response-related pathways such as the *p53 pathway, TNFα signaling via NF-κB, DNA repair, apoptosis, inflammatory response, interferon gamma response, upregulation of UV response, hypoxia, E2F targets, mitotic spindle assembly* and *G2M checkpoint* all showed similar enrichment or depletion across cell lines. Other radiation response-related pathways, such as *the reactive oxygen species pathway* and *MYC targets (version 1),* showed differential enrichment or depletion across cell lines, specifically showing enrichment only in CME037. In contrast, *EMT* was not enriched in CME037 together with depletion of *TGF-β, Notch, Wnt/β-catenin signaling,* and *angiogenesis*. Other differences include a depletion of *mTORC1* signaling and *glycolysis* in LBT005. Interestingly, we observed that the amplitude of the effect (*i.e.* fold changes) of PIR on gene expression changes was reduced in LBT005 compared to the other PDGSCLs (**Figure 2C**). This was especially evident for classical p53-dependent radiation-responsive genes (**Figure 2E**). The blunted response of LBT005 cells coincides with their elevated baseline expression of *MDM2* (**Figure 2E & Figure S1**), a gene that is amplified in this cell line (**Figure 1B**). MDM2 is a ubiquitin ligase that functions as a negative regulator of p53 by promoting its degradation. The lower induction of apoptosis-related genes such as *FDXR*, *BBC3*, *CDKN1A*, *FAS*, or *GADD45A* in LBT005 cells (**Figure S1**) may therefore partly explain their low sensitivity to PIR (**Figure 1D-E**).

**Figure 2:**
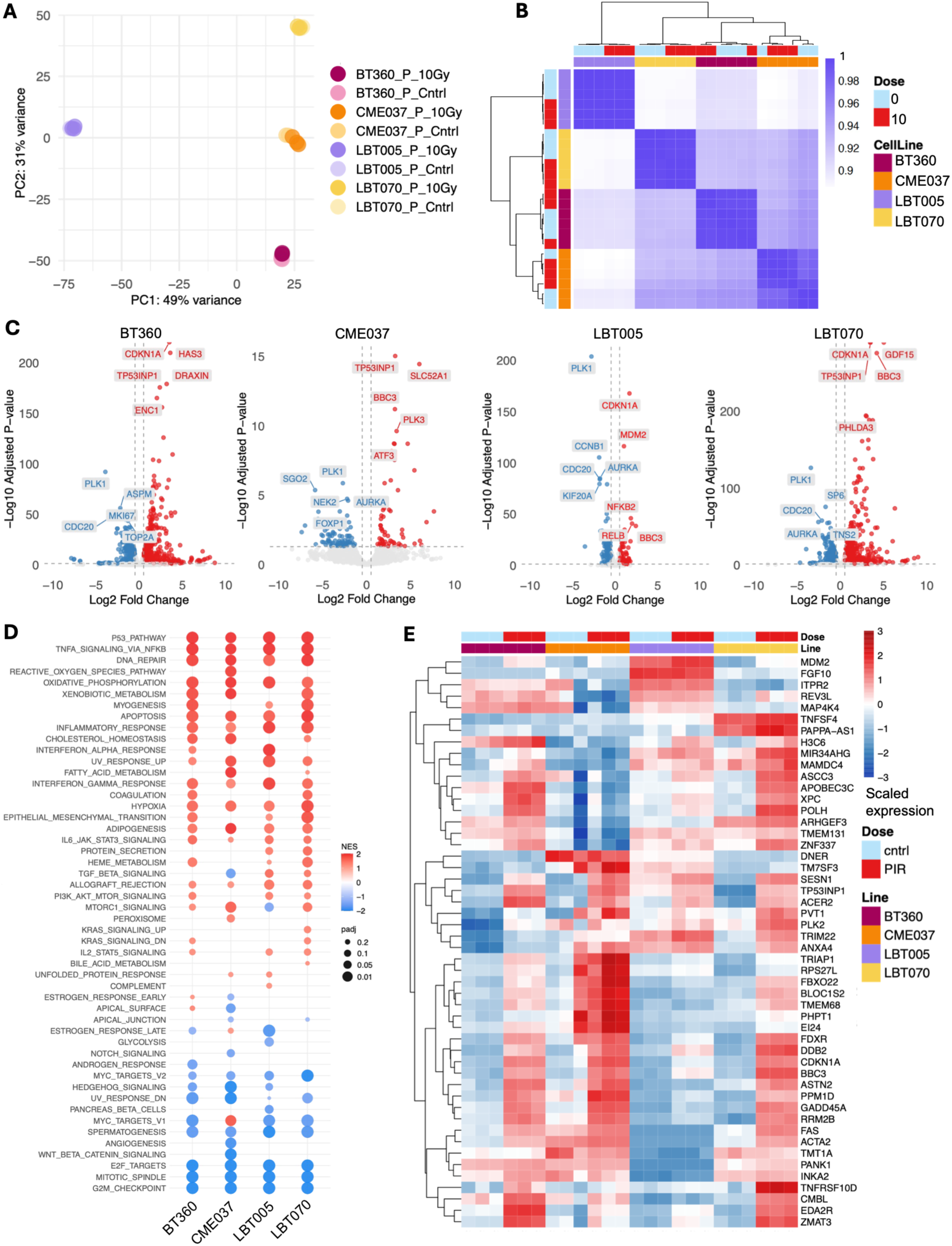
Transcriptional profile and radiation response vary between cell lines, likely driven by baseline heterogeneity. **A**: Principal component analysis (PCA) plot from bulk RNAseq data (n = 3). **B**: Sample-sample correlation plot with the top 1000 most variable genes indicating cell line as main source of variation. **C**: Volcano plots showing the DEGs for each PDGSCL after PIR. Genes that were strongly up- or downregulated (|LFC| > 0.5) and passed a significance threshold (adjusted p-value < 0.05) were highlighted in blue (downregulated) or red (upregulated). The five significant genes with highest or lowest LFC values are labeled and overall represented p53 response- and cell cycle-related genes, respectively. **D**: fGSEA showing enriched or depleted hallmark pathways per cell line after PIR. For plotting, an adjusted p-value cut-off of 0.25 was used. **E**: Scaled expression of the top 50 most variable radiation response genes across samples, based on the MSigDB gene set *MACAEVA_PBMC_RESPONSE_TO_IR*.

When comparing radiation-induced DEGs, we found both unique and overlapping sets of DEGs for each PDGSCL (**Figure 3A**). For CME037, almost all DEGs were in common with at least one of the other cell lines, and therefore, it had very few unique DEGs. For BT360, LBT005, and LBT070 55%, 57% and 47% of DEGs were common with at least one other PDGSCL. Genes that were differentially expressed after PIR in the three more responsive PDGSCLs, but not in the radioresistant line LBT005, were enriched for pathways involved in apoptosis and cellular stress responses (*e.g., FAS, DDIT4*), extracellular matrix remodeling and adhesion (*e.g., PLAU, PGF, NECTIN4*), and suppression of proliferation and cytoskeletal signaling (*e.g., SOX8, ROCK2, BCAR3, FAM72D*), suggesting that in the responsive PDGSCLs coordinated transcriptional programs are activated promoting DNA damage responses, cell death, and growth arrest following irradiation, whereas this response is largely absent in the radioresistant cell line. Enrichment analysis of the unique, PDGSCL-specific DEGs revealed cell line-specific biological processes upregulated and downregulated after PIR, except for CME037 because of the small number of unique DEGs (**Figure 3B & Table S5-6**). For BT360, several of the upregulated pathways were associated with mitochondrial respiration and oxidative phosphorylation, including *aerobic electron transport chain, ATP synthesis-coupled electron transport, respiratory electron transport chain,* and *peptidyl-serine phosphorylation,* suggesting an increase in mitochondrial metabolic activity following PIR. LBT005 was enriched for pathways associated with immune activation and antigen presentation. Consistent with this, *interferon-mediated signaling pathway* was also enriched, indicating induction of a radiation-associated inflammatory response [73]. LBT070 showed significant enrichment of pathways related to Wnt signaling, including *Wnt signaling pathway, cell–cell signaling by Wnt,* and *regulation of anoikis*.

**Figure 3:**
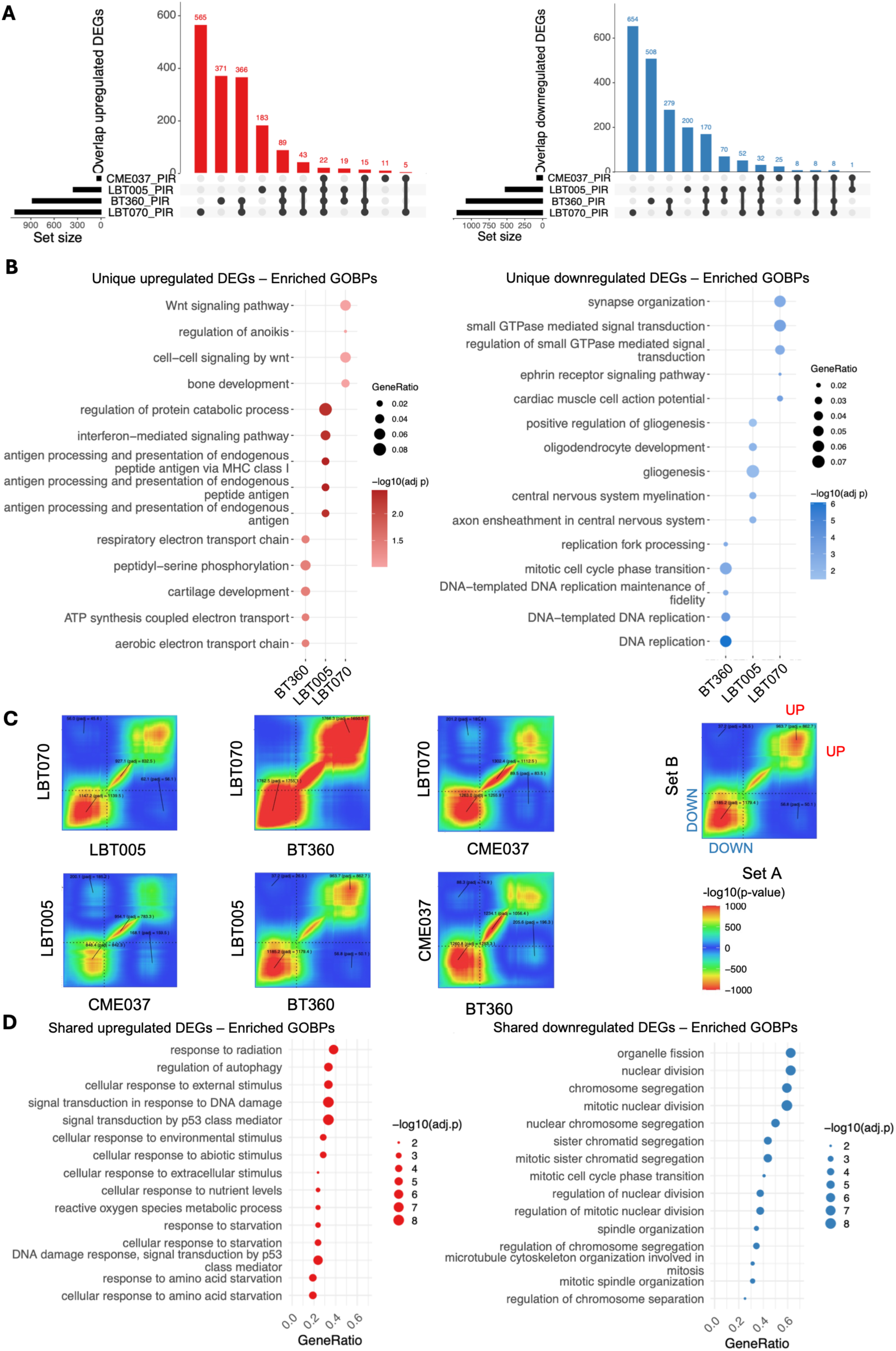
Common trends are found in the transcriptional radiation response. **A**: UpSet plots indicating unique DEGs and overlaps between upregulated (left) and downregulated (right) DEGs (|LFC| > 0) from four different PDGSCLs after PIR. **B**: Enriched biological processes (enrichGO) for the unique upregulated (left) and downregulated (right) DEGs after PIR per PDGSCL. Only significant (padj<0.25) pathways with a gene count > 4 are shown. **C**: RRHO plots comparing the 4 PDGSCLs. **D**: Enriched biological processes (enrichGO) for the shared upregulated (left) and downregulated (right) DEGs after PIR across the four PDGSCLs. For shared DEG lists, DEGs were filtered for adjusted p-value < 0.05.

Concerning unique downregulated DEGs after PIR, BT360 showed significant suppression of pathways associated with DNA replication and cell-cycle progression. Among the most significantly enriched terms were *DNA replication, DNA-templated DNA replication, DNA replication fidelity maintenance, replication fork processing,* and *the mitotic cell cycle phase transition*, indicating a broad and strong inhibition of proliferative activity following radiation exposure. In LBT005, PIR reduced the expression of pathways involved in *gliogenesis*, *oligodendrocyte development*, *central nervous system myelination*, and *axon ensheathment in the central nervous system*. These processes are all related to oligodendrocyte development and function. Given LBT005’s baseline neuronal-like phenotype, downregulation of these processes may reflect a shift away from this phenotype toward a more resistant MES-like state following irradiation [74]. Finally, in LBT070, pathways related to small GTPase-mediated signal transduction were significantly decreased. Small GTPases regulate cytoskeletal organization, cell polarity, migration, and intracellular signaling, suggesting suppression of mechanisms that are involved in GBM cell motility and proliferation [75].

Altogether, these results suggested that the intrinsic heterogeneity that exists between our PDGSCLs was reflected in their transcriptional response, and subsequently, their sensitivity to PIR.

### Despite cell line heterogeneity, common trends are found in the transcriptional response to radiation

Despite the occurrence of cell-specific transcriptional responses, the RNAseq data clearly revealed common trends in the response to radiation. Many up- and downregulated DEGs overlapped across cell lines, especially between BT360 and LBT070 (**Figure 3A**). This was confirmed by a rank-rank hypergeometric overlap (RRHO) analysis, which showed high concordance between all cell lines, but especially between BT360 and LBT070. RRHO furthermore indicated an overall more significant overlap between downregulated rather than upregulated genes (**Figure 3C**). Top up- and downregulated genes shared between multiple cell lines included *GDF15, DDB2, PLK1, CDC20, KIF14, BUB1* and *ASPM.* There were 22 upregulated DEGs shared between all four cell lines and 32 shared downregulated DEGs (**Table S3-4**). Enrichment analysis of these shared up- and downregulated DEGs indicated p53-mediated pathways, apoptosis and DNA damage signaling to be commonly upregulated and cell cycle-related processes like chromosome segregation and mitotic spindle organization as commonly downregulated, consistent with the canonical radiation response (**Figure 3D & Table S7-8**).

In addition to PIR, three PDGSCLs (BT360, LBT005 and LBT070) were irradiated with CIR because of its known higher relative biological effectiveness and its proposed potential as an alternative or additional radiation modality for GBM treatment [25, 27, 76–79]. Surprisingly, viability in response to CIR was differently influenced for each of the tested PDGSCLs. While BT360 was equally sensitive to CIR and PIR, LBT005 was more sensitive to CIR. LBT070, on the other hand, was less sensitive to CIR (**Figure S2A**). Gene expression changes in response to CIR showed strong overlap with those observed after PIR for BT360 and LBT070, and to a lesser extent for LBT005 (**Figure S2B**). In addition, we observed very similar patterns after CIR as we did after PIR, with shared downregulated DEGs being enriched for cell cycle-related processes and shared upregulated genes being enriched for p53-mediated processes (**Figure S2C-D & Table S9-10**), demonstrating that similar transcriptional mechanisms are at play early after exposure for both radiation modalities.

### Radiation consistently represses DREAM-dependent MCPH genes

To further clarify the conserved mechanisms across PDGSCLs, we focused more on the shared DEGs and the processes they are associated with. The Jensen_diseases and DisGeNET databases were used to check the expression of the shared upregulated (**Figure S3**) and downregulated (**Figure 4A**) DEGs in a disease context. Genes involved in diseases and syndromes associated with reduced head size and/or cerebral volume, such as Warburton-Anyane-Yeboa syndrome, Goldberg-Shprintzen syndrome, microcephaly and cerebral hypoplasia were strongly enriched among the downregulated DEGs. Among these were several of the MCPH genes. This corroborates previous findings indicating a p53-E2F4/DREAM-dependent repression of a subset of MCPH genes after irradiation in human cortical organoids and neural progenitor cells [45]. Given the proposed therapeutic potential of some of these MCPH genes for GBM [42], we decided to further investigate the expression of these genes in our PDGSCLs and how they were affected by irradiation.

**Figure 4:**
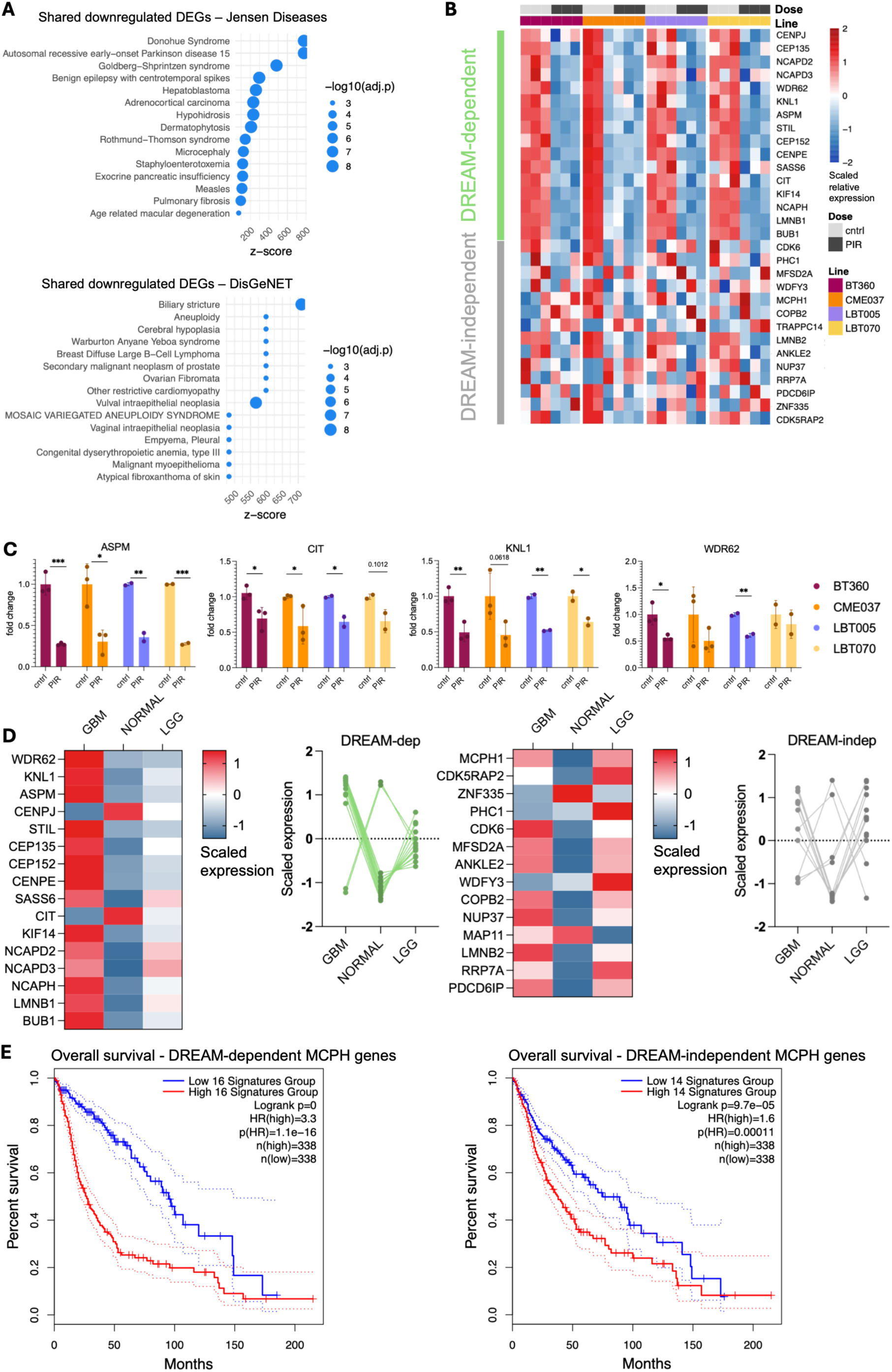
Radiation consistently represses MCPH genes. **A**: Enriched diseases from the Jensen_Diseases (top) and DisGeNET (bottom) databases among the shared downregulated DEGs after PIR. The z-score is the combined score calculated from the adjusted p-value and odds ratio to reflect both. Top enriched diseases are associated with reduced cerebral volume. **B**: Scaled relative expression of DREAM-dependent and DREAM-independent MCPH genes showing clear repression of the former after PIR. Expression of the DREAM-independent MCPH genes is more ambiguous across PDGSCLs. **C**: RT-qPCR for four DREAM-dependent MCPH genes (ASPM, CIT, KNL1, WDR62) confirms significant repression after PIR. n = 3 (BT360, CME037); n = 2 (LBT005, LBT070); error bars = SD; unpaired t-test (alpha = 0.05); * = ≤ 0.05; ** = ≤ 0.01; *** = ≤ 0.001; **** = ≤ 0.0001. **D**: Scaled expression of DREAM-dependent and DREAM-independent MCPH genes in GBM, normal tissue and low-grade glioma (LGG), showing that expression of DREAM-dependent MCPH genes is especially higher in GBM. Data from the GEPIA database. **E**: The hazard ratio for overall survival was higher for the DREAM-dependent signature (HR = 3.3; p(HR) = 1.1E-16) compared to the DREAM-independent signature (HR = 1.6; p(HR) = 0.00011). Data obtained from TCGA LGG-GBM.

First, the MCPH gene signature was divided into two subcategories, based on their DREAM-binding score from multiple chromatin immunoprecipitation experiments [46]: 16 genes with a DREAM-binding score of 7 or higher were classified as DREAM-dependent and 14 genes with a score below 7 as DREAM-independent (**Table S11**). Consistent with the known p53-dependent regulation of DREAM targets, we observed that irradiation of PDGSCLs reduced the expression of all 16 DREAM-dependent MCPH genes, whereas DREAM-independent genes were variably regulated. Genes such as *CDK6, NUP37* and *MCPH1* were upregulated after PIR in some PDGSCLs rather than being repressed (**Figure 4B**). Of the 14 DREAM-independent MCPH genes, only four showed consistent radiation-induced repression. Repression of four DREAM-dependent MCPH genes – *ASPM, CIT, KNL1* and *WDR62* – was independently confirmed with RT-qPCR (**Figure 4C**).

As for PIR, the shared downregulated DEGs after CIR were also associated with diseases characterized by reduced head size and/or cerebral volume, such as Warburton-Anyane-Yeboa syndrome, Goldberg-Shprintzen syndrome and cerebral hypoplasia (**Figure S4A)**. Looking at the MCPH genes specifically, a repression after CIR is shown as well for the majority of the DREAM-dependent MCPH genes (**Figure S4B)**. This was also suggested by RT-qPCR (**Figure S4C)**. Compared to PIR, our CIR samples seemed to carry more variation between replicates, making the repression of DREAM-dependent MCPH genes less apparent. For CIR, only two replicates were available, affecting the statistical significance.

Nonetheless, these results suggest that repression of DREAM-dependent MCPH genes is independent of radiation quality.

### MCPH genes are overexpressed in GSCs and correlate with poor prognosis

To highlight the potential importance of the MCPH genes in glioma biology, we analyzed their expression using the Gene Expression Profiling Interactive Analysis (GEPIA) database [69]. Most DREAM-dependent MCPH genes are strongly enriched in GBM and, to a lesser extent, in low-grade glioma (LGG) compared to normal brain tissue. *CENPJ* and *CIT*, however, have an inverse expression profile. In contrast, the expression of DREAM-independent MCPH genes is more similar between GBM and LGG but remains lower overall in normal tissue (**Figure 4D**). The low expression of MCPH genes in adult normal brain tissue aligns with their proposed importance for the function of NPCs, which are only sparsely present in the adult brain. This is also reflected by the developmental expression of MCPH genes in brain tissue, which again shows some discrepancy between DREAM-dependent and DREAM-independent genes. While DREAM-dependent MCPH genes (except for *CIT*) are especially highly expressed during the earliest stages of gestation and neurogenesis, most DREAM-independent MCPH genes retain expression until late gestation, and some even postnatally (**Figure S5**). Early stages of brain development are characterized by a high prevalence of NPCs, such as radial glia, that resemble GSCs [28].

Expression of MCPH genes, particularly DREAM-dependent, was enriched in GSCs, but not GBM tumor cells (non-GSCs) and in recurrent GBM compared to astrocytoma, newly diagnosed GBM, and oligodendroglioma from single-cell RNAseq datasets of clinical samples [7, 80] (**Figure S6**). Furthermore, based on gene expression profiles in glioma patient cohorts from TCGA LGG-GBM, we found that expression of both DREAM-dependent and -independent MCPH gene signatures were associated with poorer disease outcome (**Figure 4E**). The hazard ratio for overall survival was, however, higher for the DREAM-dependent signature (HR = 3.3; p(HR) = 1.1E-16) compared to the DREAM-independent signature (HR = 1.6; p(HR) = 0.00011), suggesting it is associated with a poorer prognosis.

### Repression of MCPH genes is associated with reduced expression of FOXM1

To investigate potential regulatory mechanisms of radiation response, we analyzed transcription factor (TF) regulatory network activity and binding after PIR. We observed increased TP53 activity across all cell lines. Other TFs with increased activity in at least two out of four cell lines included HIF1A and NFE2L2. Decreased activity of E2F2 and FOXM1 was observed across all cell lines, and other TFs with reduced activity in at least two out of four cell lines included E2F1, E2F2, E2F3, E2F4, E2F5, DUX4 and ZFX (**Figure 5A**).

**Figure 5:**
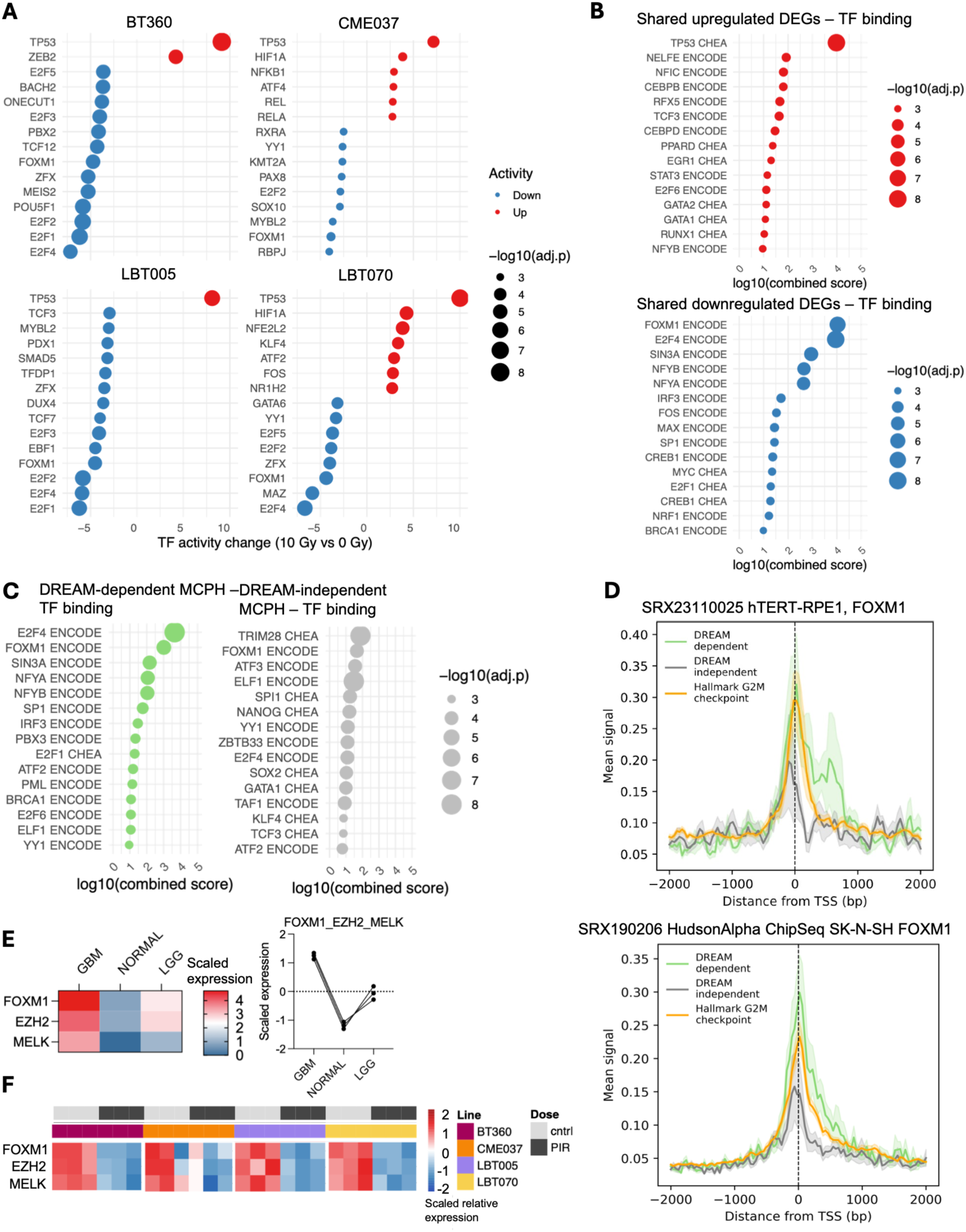
Radiation-induced repression of MCPH genes is associated with reduced activity and expression of *FOXM1*. **A**: Transcription factor (TF) activity (Dorothea & VIPER) after irradiation per PDGSCL. Activity of TP53 is increased in all 4 cell lines, while activity of FOXM1 and E2F2 is decreased in all cell lines after PIR. **B**: TF binding enrichment in the shared upregulated and downregulated DEGs in all cell lines (ENCODE & ChEA Consensus), suggesting TP53 and FOXM1 as top regulators for upregulated (top) and downregulated (bottom) genes, respectively. **C**: TF binding enrichment for the DREAM-dependent (top) and -independent (bottom) MCPH genes, suggesting FOXM1 as a regulator of both sets, but with a higher combined score (*i.e.* combination of odds ratio and adjusted p-value) for DREAM-dependent MCPH genes. **D**: ChIP-seq signal scoring over gene signature TSS regions for FOXM1. Data obtained from ChIP-Atlas. **E**: *FOXM1*, *EZH2* and *MELK* are overexpressed in GBM compared to normal tissue and low-grade glioma (LGG). Data from the GEPIA database. **F**: Radiation decreases the expression of *FOXM1, EZH2* and *MELK* in all cell lines.

The shared upregulated DEGs across all cell lines were predominantly enriched for TP53 targets (**Figure 5B**), consistent with its increased activity after PIR. Considering the shared downregulated DEGs, we observed a strong enrichment for FOXM1 targets, also consistent with its decreased activity. Another important regulator of commonly downregulated genes was E2F4, a known repressor of cell cycle-related genes and a central component of the DREAM complex [81, 82].

Considering the top regulators of the MCPH genes, a large overlap existed with the regulators of the shared downregulated DEGs and those of the DREAM-dependent MCPH genes, including E2F4, FOXM1, SIN3A, NFYA, NFYB, and IRF. Interestingly, DREAM-dependent and, albeit to a lesser extent, DREAM-independent MCPH genes were both enriched for FOXM1 targets (**Figure 5C**). FOXM1 is a regulator of cell cycle genes, especially those involved in the G2M phase [83]. In the context of cancer, *FOXM1* is often co-expressed with *MELK* and has been shown to target *EZH2* to promote progression and radioresistance [47, 84–87]. Metagene profiles of FOXM1 ChIP-seq signal centered on transcription start sites (TSS) were generated based on ChIP-Atlas [71] and stratified by functional gene sets including DREAM-dependent MCPH genes, DREAM-independent MCPH genes, and Hallmark G2M checkpoint as canonical FOXM1 targets (**Figure 5D**). In two human cell lines, hTERT-immortalized retinal pigment epithelial cells and neuroblastoma cells, DREAM-dependent and the hallmark G2M checkpoint gene sets displayed strong enrichment peaks at the TSS, while the DREAM-independent gene set displayed a lower peak, suggesting a stronger association of FOXM1 with the promoters of DREAM-dependent compared to DREAM-independent MCPH genes.

Interestingly, in GBM, *FOXM1, EZH2,* and *MELK* are overexpressed relative to normal tissue and low-grade glioma (**Figure 5E**), resembling the expression patterns of most DREAM-dependent MCPH genes in these contexts (**Figure 4D**). A strong correlation between expression profiles of DREAM-dependent MCPH genes (except *CIT*) and those of FOXM1, EZH2 and MELK was also found using glioma data from the Chinese Glioma Genome Atlas (CGGA), whereas these correlations were overall less significant for DREAM-independent MCPH genes (**Figure S7**). Together, these results suggested (1) that radiation repressed *FOXM1* in PDGSCLs, and (2) that FOXM1 regulates the expression of DREAM-dependent MCPH genes.

### Radiation-induced repression of MCPH genes is heterogeneous and likely caused by GBM cell state redistribution

GBM is characterized by extensive intratumoral heterogeneity, as is reflected by the heterogeneous composition of the PDGSCLs used in this study (**Figure 1C**). In addition to our observation of a consistent repression of MCPH genes across these cell lines, we aimed to address the intratumoral heterogeneity with RNA sequencing at a single-cell resolution. For this, CME037 cells were irradiated with 10 Gy and compared to sham-irradiated cells with scRNAseq after 6 hours. UMAP clustering indicated a shift in transcriptional profiles after irradiation (**Figure 6A**). Given the roles of MCPH genes in different cell cycle-related processes, S and G2M scores were regressed out to minimize the confounding effects of cell cycle state, thereby facilitating the identification of treatment-induced transcriptional changes and biologically relevant cellular heterogeneity. These regressed data were used for downstream analysis. A DREAM-dependent and -independent module score was calculated for each cell, revealing that cells with high DREAM-independent scores were more uniformly distributed across conditions, while control cells were enriched for high DREAM-dependent scores, again confirming radiation-induced repression of (mostly DREAM-dependent) MCPH genes (**Figure 6B**). We therefore continued our analysis focusing on the DREAM-dependent signature. Reduced expression of top-hit transcription factors *E2F4* and *FOXM1* (**Figure 5B-C**), together with FOXM1 targets *EZH2* and *MELK,* was additionally confirmed post-irradiation (**Figure 6C**).

**Figure 6:**
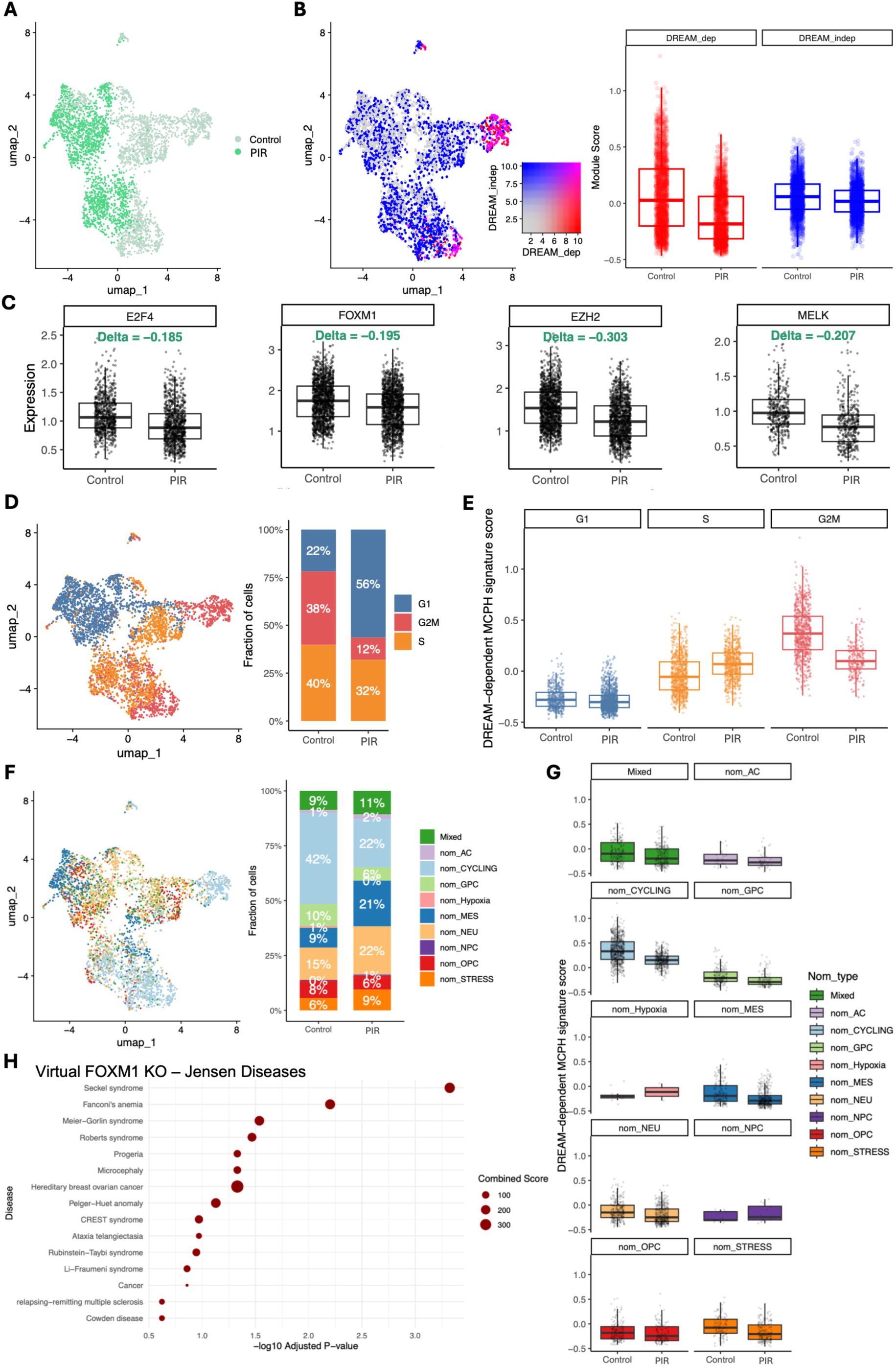
scRNAseq suggests radiation-induced repression of MCPH genes is linked to redistribution of GBM cell phenotypes. **A**: UMAP showing distinction between (sham-) irradiated cells (PIR 10 Gy); data were randomly downsampled to 2000 cells/condition, and cell cycle effects were regressed out. **B**: Cells with high DREAM-independent score are scattered across conditions, whereas cells with high DREAM-dependent score are enriched in control and depleted after PIR. **C**: Expression of top hit TFs E2F4 and FOXM1, together with FOXM1 targets EZH2 and MELK, is reduced after PIR. Delta scores indicate directionality of change in expression, with negative scores indicating reduced expression. **D**: PIR results in redistribution of cell cycle phases, namely an enrichment of the G1 phase and a depletion of the G2/M phase. **E:** DREAM-dependent MCPH signature score across cell cycles. PIR results in depletion of cells with high DREAM-dependent MCPH signature scores, here G2/M phase cells. **F**: PIR results in redistribution of Nomura GBM phenotypes, namely an enrichment of MES- and NEU-like cells and a depletion of cycling cells. **G**: DREAM-dependent MCPH signature score across phenotypes. PIR results in depletion of cells with high DREAM-dependent MCPH signature scores, here, cycling cells. **H**: Jensen disease enrichment after virtual FOXM1 KO, suggesting a strong causality between loss of FOXM1 and diseases associated with reduced head size and/or brain volume, including microcephaly, thereby strengthening the regulatory role of FOXM1 in MCPH gene expression.

Next, cells were stratified based on cell cycle phases. Irradiation led to a marked enrichment of cells in G1 and a concomitant depletion of G2/M cells (**Figure 6D**). When stratifying cells by cell cycle phase, the lowest DREAM-dependent module scores were observed in G1-phase cells and the highest scores in G2/M-phase cells (**Figure 6E**). This suggests that radiation shifts the cell population toward a cell cycle state (G1) in which the signature is intrinsically low, while depleting states (G2) in which the signature score is higher.

Next, cells were assigned to one of the Nomura GBM cell states [11]. Most clearly, radiation induces an increase in MES- and NEU-like cells while reducing the cycling cell population (**Figure 6F**). Cells assigned to the cycling phenotype can have this phenotype on top of another neurodevelopmental-like phenotype, as it merely emphasizes their proliferative capacity [11]. The original publication did not consider this phenotype as a separate, exclusive state, but given the importance of the cell cycle in this study, we decided to include it in our analysis as a separate state. This again suggests that radiation depletes a phenotype with an intrinsically high DREAM-dependent MCPH score while shifting to phenotypes with intrinsically lower scores (**Figure 6G**). Therefore, population-level downregulation likely reflects radiation-induced redistribution of cell cycle states and phenotypes rather than uniform per-cell repression.

Finally, to verify the causal relationship between reduced *FOXM1* expression and repression of MCPH genes, a virtual *FOXM1* knockout was performed on these single-cell data. We used scTeniFoldKnk, which allows us to simulate the virtual knockout of a gene in single-cell gene regulatory networks to identify which other genes and biological programs are most affected by its loss [63]. Although it cannot provide actual changes in mRNA expression after perturbation, it can estimate regulatory dependencies. Enrichment analysis for biological processes after virtual *FOXM1* knockout indicated significant enrichments for cell cycle-related processes, suggesting FOXM1 predominantly affects genes enriched for cell-cycle and mitotic programs and has strong upstream regulatory control of proliferative networks (**Figure S8**). Virtual *FOXM1* knockout also resulted in top significant enrichments for diseases associated with reduced head size and/or brain volume, including microcephaly, Seckel syndrome, Fanconi anemia, Meier-Gorlin syndrome, Roberts syndrome and progeria (**Figure 6H**), providing additional arguments for the regulatory role of FOXM1 in MCPH gene expression.

### Pharmacological FOXM1 inhibition represses MCPH genes and enhances radiosensitivity

We sought to verify the causal link between reduced FOXM1 expression, activity and repression of MCPH genes, and to investigate the potential therapeutic value of inhibiting FOXM1 and MCPH gene expression. We continued our analysis with the U251-MG cell line due to its rapid growth and availability. These cells carry a p53 mutation (missense point mutation R273H, gain-of-function). Significant radiation-induced repression of *ASPM* and *KNL1* was observed in this cell line (**Figure 7A**). Radiation-induced repression of CIT and WDR62 was less apparent and not significant. *FOXM1* expression did not significantly decrease after PIR. Thus, as a proxy for reduced FOXM1 activity after irradiation in U251 cells, we measured the expression of one of its canonical targets and activator, PLK1 [88, 89], which was also among the top downregulated genes across all four PDGSCLs (**Figure 2C**). In U251 cells, *PLK1* expression was also significantly reduced (**Figure 7A**). Because PLK1 is essential for FOXM1 activity, its strong downregulation might lead to a significant reduction in FOXM1’s transcriptional effect without affecting FOXM1’s mRNA levels.

**Figure 7:**
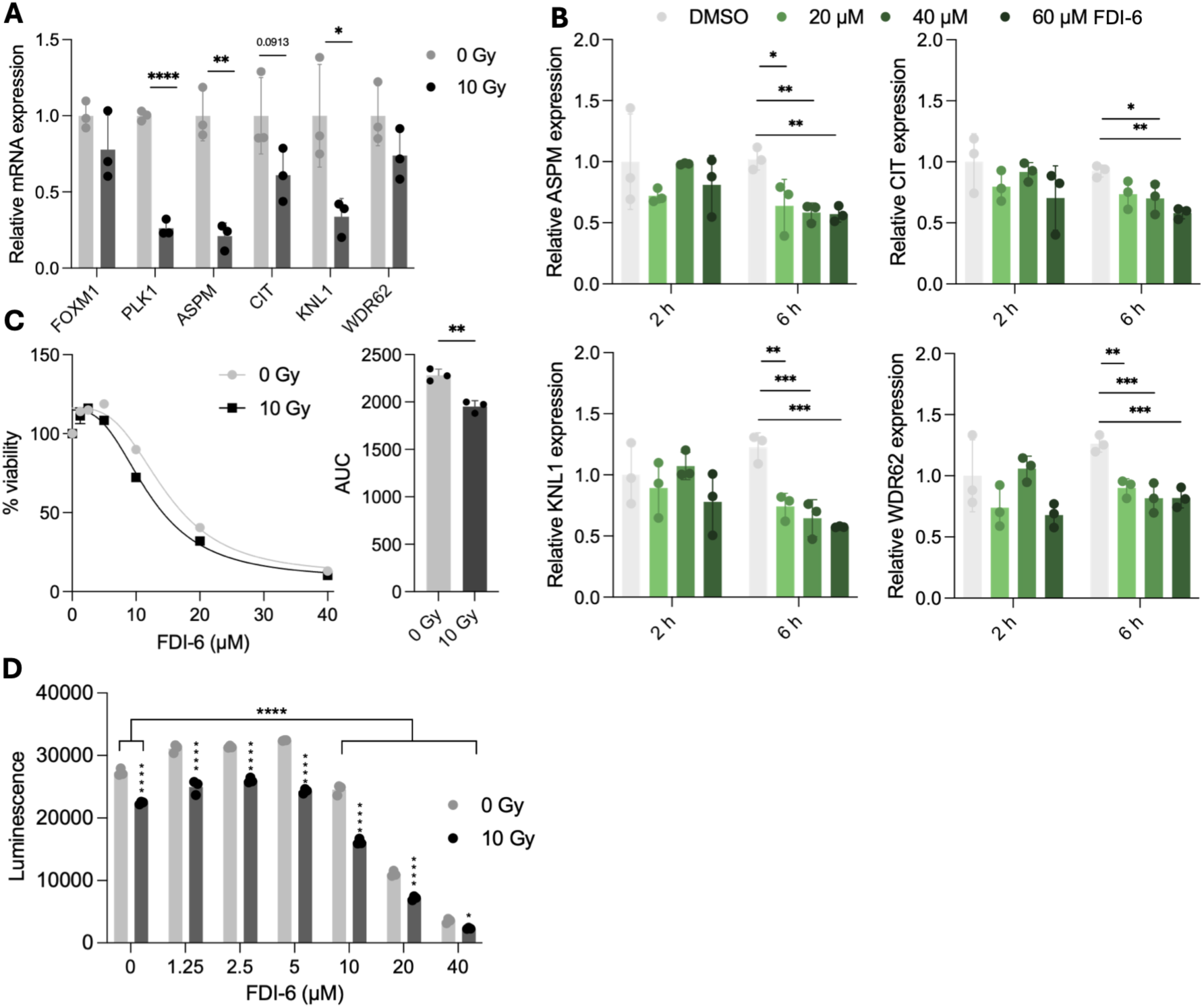
Pharmacological inhibition of *FOXM1* reduces expression of MCPH genes and viability in U251 cells. **A**: RT-qPCR for *FOXM1, PLK1,* and DREAM-dependent MCPH genes (*ASPM, CIT, KNL1, WDR62*) in p53-mutant U251 cells confirms significant repression of *PLK1, ASPM* and *KNL1* after PIR. n = 3; error bars = SD; unpaired t-test (alpha = 0.05); * = ≤ 0.05; ** = ≤ 0.01; *** = ≤ 0.001; **** = ≤ 0.0001 **B**: Treatment with FOXM1 inhibitor FDI-6 resulted in repression of MCPH genes *ASPM*, *CIT*, *KNL1, and WDR62* after 6 but not after 2 hours. n = 3; error bars = SD; Ordinary One-way ANOVA with Šídák’s multiple comparisons test (alpha = 0.05); * = ≤ 0.05; ** = ≤ 0.01; *** = ≤ 0.001; **** = ≤ 0.0001. **C**: Dose-response curve (non-linear regression) for increasing concentrations of FDI-6 in control and irradiated cells. 0 µM FDI-6 = 100% viability for both control and irradiated cells. AUC values indicate a significant added effect of PIR. n = 3; error bars = SD; Unpaired t-test (alpha = 0.05); * = ≤ 0.05; ** = ≤ 0.01; *** = ≤ 0.001; **** = ≤ 0.0001. **D**: 10, 20 and 40 µM of FDI-6 and 10 Gy PIR significantly reduced ATP-based viability. n = 3; error bars = SD; 2-way ANOVA with Šídák’s multiple comparisons test (alpha = 0.05); * = ≤ 0.05; ** = ≤ 0.01; *** = ≤ 0.001; **** = ≤ 0.0001.

The effect of FOXM1 inhibition on MCPH gene expression was investigated using a small-molecule inhibitor of FOXM1, FDI-6. FDI-6 has been shown to bind FOXM1 and thereby cause downregulation of its targets in breast cancer cells [90] and has since been used in a preclinical setting in multiple cancer studies [91–93]. No significant changes in MCPH gene expression were observed after 2 h of FDI-6 treatment, whereas significant repression of all investigated MCPH genes was observed after 6 h across all tested concentrations (**Figure 7B**). These findings are consistent with a regulatory role of FOXM1 in MCPH gene expression independent of p53 status. In addition, FDI-6 dose-response was calculated for both irradiated and non-irradiated cells (**Figure 7C-D**). Non-linear regression curves and Area Under the Curve (AUC) values indicate a significant added effect of PIR at intermediate doses of FDI-6, suggesting inhibition of FOXM1 6 hours before PIR increases radiosensitivity of U251 cells.

Overall, these results supported a functional association between FOXM1 activity and MCPH gene expression in GBM cells. Pharmacological inhibition of FOXM1 further indicated a regulatory role for FOXM1 in selected MCPH genes and FOXM1 inhibition prior to PIR enhanced radiosensitivity, suggesting that FOXM1 acts upstream to coordinate a broader proliferative program that contributes to radiation response.

## Discussion

With its vast inter- and intratumoral heterogeneity, high level of resistance to standard-of-care treatment and poor prognosis, GBM poses a great clinical challenge. Many attempts have been made in the past decades to overcome this challenge; unfortunately, without much success. Here, we investigated the response of four different PDGSCLs across the spectrum of GBM subtypes to low- and high-LET irradiation to potentially uncover novel vulnerabilities.

### Cell line heterogeneity drives sensitivity and transcriptional response to PIR

The transcriptional response to radiation can vary between molecular subtypes, contributing to heterogeneous growth arrest and survival. For example, radiation induces robust cell-cycle arrest in proneural glioma models, characterized by strong p53 activation and loss of proliferative activity. In contrast, radiation can promote a transition toward a mesenchymal-like state associated with activation of inflammatory signaling pathways, including STAT3 and C/EBPβ, which are linked to therapy resistance [94–96].

Our most sensitive line, BT360, is most strongly negatively enriched for mesenchymal-associated signatures, agreeing with the concept of mesenchymal-like cells being more radioresistant. In contrast, the most resistant line, LBT005, does show enrichment of the RICHARDS_INJURY signature, reflecting an inflammatory-like phenotype, although it does not show strong enrichment of other mesenchymal-like signatures. Instead, neuronal-like signatures are enriched for this cell line. Although generally, mesenchymal-like states are often linked to high treatment resistance [95], recent studies have put forward neuronal-like states to drive GBM progression, linking their invasion to developmental neuronal migration and indicating their high level of stemness and plasticity as determining features of GBM recurrence [10, 23, 97, 98]. This confirms the complex relationship between GBM cell state and radiation response. Across PDGSCLs, irradiation induced a differential enrichment of response pathways and expression of canonical radiation response genes (**Figure 2D-E**). Here, the least responsive and *MDM2*-amplified PDGSCL, LBT005, indeed showed a limited induction of apoptosis-related genes compared to the more sensitive PDGSCLs. By targeting p53 for degradation, amplified *MDM2* can lead to suppression of radiation-induced cell cycle arrest, senescence, and apoptosis, linking it to radioresistance [99]. Thus, the amplitude of the p53-mediated response of apoptosis-related genes may be a proxy for radiation sensitivity.

The unique, PDGSCL-specific responses could be associated with markedly different radiation sensitivities, suggesting that specific stress-response programs may influence radiosensitivity. The most radiosensitive cell line, BT360, displayed upregulation of oxidative phosphorylation and mitochondrial respiratory pathways together with suppression of DNA replication and cell-cycle progression. Suppression of proliferative programs suggests that these cells may undergo a relatively more effective radiation response characterized by cell-cycle arrest and reduced replication, potentially explaining their high radiation sensitivity. Increased oxidative phosphorylation has been associated with stemness and radioresistance [100]. In contrast, it has also been reported that it can lead to increased production of mitochondrial reactive oxygen species and oxidative damage in irradiated cells, thereby contributing to radiosensitivity [101]. LBT005 exhibited activation of interferon signaling, antigen presentation, and other immunogenic pathways, together with suppression of gliogenesis and oligodendrocyte differentiation-associated programs. This observation is consistent with LBT005’s baseline enrichments of neuronal-like and injury response signatures. The activation of immunogenic, injury response-like pathways with the concomitant loss of differentiation-associated transcriptional programs may indicate a shift towards a more stem-like and stress-adapted phenotype. Such dedifferentiation-like transitions have previously been linked to treatment resistance in GBM, where treatment can drive cells away from lineage-restricted states and toward more stem-like or injury-responsive cellular states [7, 28]. Therefore, although irradiation seemingly increased immunogenic signaling, the overall transcriptional response may represent an adaptive state that facilitates tumor cell survival. Taken together, these findings suggest that radiation response in GBM is associated not only with induction of the canonical DNA damage response, but also with specific adaptive transcriptional states that emerge shortly after irradiation (6 hours) [94]. Additionally, they suggest that intrinsic phenotypes can determine transcriptional radiation response and radiosensitivity, thereby highlighting the detrimental effect of heterogeneity between GBM patients.

Despite these cell-specific responses, shared radiation-induced biological processes were evident across PDGSCLs (**Figure 3C-D & Table S3-4**). Conforming to the canonical radiation response, an upregulation of p53-related pathways coincided with a downregulation of cell cycle processes. The same was observed for CIR, even though LBT005 and LBT070 were not equally resistant or sensitive to CIR as they were to PIR, respectively.

### Radiation consistently represses a subset of MCPH genes that are specifically associated with stemness, glioma grade and prognosis

Our bulk RNAseq data indicate a global shutdown of proliferation-associated transcription after PIR. Among these cell cycle-related genes, we highlight a particular set of genes linked to microcephaly. Some of these genes are well-established components of the mitotic cell cycle machinery [43], and previous research has identified the regulatory role of the p53-E2F4/DREAM axis in radiation-induced repression of these genes in human neural progenitor cells [45]. In accordance with this, DREAM-dependent MCPH genes were coordinately downregulated in all PDGSCLs following PIR (**Figure 4B**). Our scRNAseq data, however, suggest that the observed repression of MCPH genes at the bulk level does not necessarily reflect a per-cell repression but rather shifts in the composition of the cell population. Here, phenotypes reflecting cells with high cell cycle activity were depleted, consistent with the involvement of the MCPH genes in this process. Cell state transitions have been increasingly recognized as a key component of therapy response in GBM and are known to contribute to treatment resistance and recurrence [23, 95].

The developmental function of MCPH genes provides a compelling rationale for their investigation in GBM. Increasing evidence indicates that GBM recapitulates key aspects of neurodevelopment and GSCs exhibit transcriptional and functional similarities to NPCs and radial glia-like progenitors [10, 28, 29]. Therefore, the response to radiation is likely very similar between NPCs and GSCs [29, 45]. This resemblance raises the possibility that genes required for NPC proliferation may also represent critical dependencies in GBM, particularly during recovery from radiation-induced damage. Interestingly, Zika virus (ZIKV), which preferentially infects NPCs and is responsible for the congenital microcephaly phenotype, has emerged as a promising oncolytic agent against GBM due to its selective tropism for GSCs via a SOX2–integrin αvβ5 axis [102]. This further supports the concept that developmental pathways active during neurogenesis are reactivated in GSCs and may represent therapeutically exploitable vulnerabilities.

Among MCPH genes, several stand out as particularly attractive therapeutic targets. *ASPM* is consistently overexpressed in high-grade gliomas and has been implicated in stem-cell maintenance, mitotic progression, and tumor growth [54, 103, 104]. Similarly, *KNL1 [105], CENPE [106], and KIF14 [106, 107],* have been shown to regulate processes that are indispensable for the rapid proliferation characteristics of high-grade glioma and GBM cells [42]. In addition, some MCPH genes are included in the *Cycling* metaprogram defined by Nomura *et al.* (2025) for GBM, including *ASPM, CIT, KNL1, CENPE, LMNB1, KIF14 and STIL* [11]. This suggests that these genes are involved in the proliferative compartment of GBM and are likely most highly expressed in actively dividing malignant cells, and particularly progenitor-like tumor populations. Moreover, Mathur *et al*. (2024) investigated tumor-wide GBM features and heterogeneity. One of their modules, *R_turquoise*, is characterized by neurodevelopmental programs and correlates with a high sample purity, suggesting these mechanisms act in malignant cells across all patients in their cohort. This module was enriched for DREAM-dependent MCPH genes *WDR62, KNL1, ASPM, STIL, CENPE, SASS6, KIF14, NCAPD2, NCAPD3, NCAPH, LMNB1* and *BUB1,* and for DREAM-independent MCPH genes *MCPH1, ANKLE2* and *LMNB2* (**Table S11**), suggesting they are associated with malignant cells and may contribute to the maintenance of the proliferative intermediate progenitor-like state identified by Mathur *et al.* Rather than representing region-specific adaptations, these genes appear to participate in a tumor-wide proliferative program, making them attractive candidates for further investigation as therapeutic targets [108].

While their loss during embryogenesis critically disrupts cortical expansion, accumulating evidence suggests that several MCPH genes become largely dispensable in the postnatal organism. Indeed, many neurodevelopmental programs that lie at the crossroads between cancer and neurodevelopment have limited functional relevance after completion of cortical development (**Figure S4)** [30]. In contrast, proliferating tumor cells remain critically dependent on these pathways, potentially creating a vulnerability that can be therapeutically exploited.

Taken together, the particularly high expression of DREAM-dependent MCPH genes during early brain development, and in GSCs compared to more differentiated GBM cells, suggests they are related to stemness. Furthermore, the higher expression in GBM compared to LGG, in recurrent compared to primary GBM, and the lower survival among patients with high expression indicate the clinical relevance of this gene signature. This suggests that these genes are tumorigenic and may constitute therapeutic targets.

Targeting MCPH-associated pathways may also offer advantages over conventional microtubule-targeting agents (MTAs). Such drugs as vinca alkaloids and taxanes effectively disrupt mitosis but are associated with severe systemic toxicities, including neurotoxicity, myelosuppression, and limited brain penetration [41, 42]. In contrast, inhibition of MCPH genes could achieve selective disruption of mitotic machinery in GBM cells while sparing the majority of differentiated adult tissues [42]. Despite promising preclinical studies, targeting individual MCPH genes may be insufficient because of partial functional redundancy and the ability of cancer cells to use alternative mitotic networks [109, 110]. An alternative strategy could be to identify common upstream regulators of MCPH gene expression. Such regulators may provide broader therapeutic effects than inhibition of a single downstream effector. Identification of these shared regulatory mechanisms could reveal novel opportunities to simultaneously suppress multiple MCPH-dependent pathways and enhance radiosensitivity.

### MCPH gene expression is regulated by FOXM1, whose functional inhibition leads to enhanced radiosensitivity

Beyond DREAM-dependency, we found that MCPH genes are particularly enriched among FOXM1 targets, and FOXM1 binds to their promoters in several neuronal cell types. Furthermore, FOXM1 is one of the most significantly downregulated genes after PIR in all PDGSCLs. FOXM1 is a master transcriptional regulator of cell-cycle progression, mitosis, DNA replication, and DNA damage repair, and is known to interact with the DREAM complex to regulate cell cycle-dependent gene expression [47, 111]. Accordingly, we show that expression of the DREAM-dependent MCPH gene subset is highly correlated with FOXM1 across multiple datasets.

FOXM1 has been extensively implicated in resistance to radiotherapy through multiple complementary pathways [48, 112, 113]. *FOXM1* expression is elevated in GBM relative to normal brain tissue (**Figure 5E**) and correlates with tumor growth, stem-cell signatures, and poor clinical outcome in solid tumors [49, 50, 112]. Substantial preclinical evidence supports therapeutic targeting of FOXM1 in GBM. Inhibition of FOXM1 reduces proliferation, self-renewal, invasion, and tumorigenicity while sensitizing glioma cells to radiation and temozolomide [49, 51, 113]. Despite these encouraging findings, clinical translation of FOXM1 inhibition remains challenging. TFs are generally considered very difficult to target, mainly due to their structural inaccessibility [114]. Available FOXM1 inhibitors often exhibit limited specificity, suboptimal pharmacokinetic properties, and uncertain blood-brain barrier penetration. An alternative strategy for direct TF targeting is the use of proteolysis-targeting chimeras (PROTACs). These selectively bind the TF of interest using a specific protein ligand and target the TF for degradation by recruiting an E3 ubiquitin ligase [115]. FOXM1 PROTACs have been tested preclinically in cancer cells and xenografts and showed promising results, including reduced cancer cell proliferation and tumor growth [116, 117].

Because FOXM1 regulates a broad range of proliferative pathways, systemic inhibition may result in toxicity in normal actively dividing tissue, for example, during wound healing. Resting, non-proliferative tissue is, therefore, thought to be less severely affected. *In vivo* studies have indeed reported anti-tumor effects of FOXM1 inhibition, while severe side effects remained absent [118], again highlighting the therapeutic benefit of FOXM1 inhibition over MTAs.

The involvement of FOXM1 in cell-cycle regulation, DNA repair, stem-cell maintenance, and MCPH gene expression make it a potential candidate for therapeutic intervention. Rather than targeting individual MCPH genes, inhibition of FOXM1 may provide a means of simultaneously suppressing multiple gliomagenesis- and radioresistance-associated programs that are (re)activated in GBM, thereby potentially enhancing the efficacy of radiotherapy while limiting compensatory pathway activation.

Pharmacological inhibition of FOXM1 resulted in significant downregulation of *ASPM, CIT, KNL1,* and *WDR62* after 6 hours, but not after 2 hours, across all tested concentrations. Importantly, this effect was observed in U251 cells, which harbor a p53 missense mutation (R273H), indicating that FOXM1-dependent regulation of MCPH genes may occur independently of p53 status. However, *FOXM1* inhibition did increase radiosensitivity (**Figure 7C-D**).

The repression of MCPH genes following FDI-6 treatment is consistent with FOXM1 contributing to their regulation, although indirect and FOXM1-independent effects cannot be excluded. Together with the enhanced radiation response, these findings provide additional preclinical evidence supporting FOXM1 inhibition as a radiosensitizing strategy in GBM.

Taken together, our findings support a model in which PIR induces both differential responses that reflect radiosensitivity and a coordinated collapse of the mitotic transcriptional program in GBM cells through both direct transcriptional repression and changes in cellular composition. This process involves reduced FOXM1 activity and a shift away from proliferative cell states. These mechanisms are likely not mutually exclusive but instead act in concert to enforce a robust anti-proliferative response to DNA damage. We particularly highlight the radiation-induced repression of DREAM-dependent MCPH genes associated with reduced FOXM1 activity. By unravelling the upstream transcriptional regulation of the MCPH genes as a whole, we hope to provide insights into the collective targeting of multiple MCPH genes, potentially yielding more promising results. This way, acute inhibition of FOXM1 function may enhance the efficacy of PIR by reinforcing the shutdown of proliferative programs.

### Limitations and future perspectives

Our analyses are primarily based on a small cohort of PDGSCLs using transcriptional data. Future studies incorporating larger cohorts, protein-level measurements, and more functional readouts at later time points would provide further mechanistic insight. Moreover, mechanistic experiments were performed using U251 cells rather than PDGSCLs, due to practical constraints, while we acknowledge that the use of PDGSCLs would be more relevant to link to our bulk and single-cell RNAseq data. Another potential limitation of our approach is that FDI-6 exerts biological effects beyond FOXM1 inhibition. Although originally characterized as a selective inhibitor of FOXM1 DNA binding, more recent studies have demonstrated that FDI-6 also activates the aryl hydrocarbon receptor, suggesting that part of its anti-proliferative or radiosensitizing activity may result from FOXM1-independent mechanisms [119]. We acknowledge that targeting TFs is particularly challenging, fueling the search for alternative strategies, such as PROTACs and phytochemicals [120, 121]. Additionally, our findings are derived from *in vitro* models, and validation in *in vivo* systems will be necessary to assess clinical relevance. Finally, our CIR data were limited by a small sample cohort and further investigation of CIR is warranted.

Despite these limitations, our study provides important insights into the transcriptional response of GBM cells to RT. The identification of FOXM1 as a potential regulator of MCPH gene expression highlights a potential vulnerability that could be exploited therapeutically. Finally, our single-cell findings underscore the importance of considering both transcriptional regulation and cell-state dynamics when interpreting therapy-induced changes in heterogeneous tumor systems.

## Data availability

RNAseq data generated in this study are available at ArrayExpress E-MTAB-17533. Any additional information for reanalysis is available from the corresponding author upon reasonable request.

## Statements & Declarations

## Acknowledgements

We would like to thank Jo Belmans, Marijke Jaf, Ann Janssen, Yanti De Visser, Basiel Cole, and Niels Vandamme for technical assistance. BT360 cells were kindly provided by the Dana-Farber Cancer Institute (Boston, MA, USA).

## Author contributions

Conceptualization: R.Q., F.D.S., K.T., I.P., and L.V.E.; methodology: R.Q., E.E., K.T., and L.V.E.; investigation: R.Q., L.V.E.; data curation: E.E.; formal analysis: E.E., L.V.E. and R.Q.; visualization: L.V.E., E.E., and R.Q.; supervision: R.Q., I.P., and F.D.S.; writing - original draft: L.V.E. and R.Q.; writing - review and editing: A.P-K., K.T., I.P., and F.D.S. All authors have read and agreed to the published version of the manuscript.

## Funding

The results from the CIR experiments presented here are based on Experiment SBio-08 performed at the GSI Helmholtzzentrum für Schwerionenforschung, Darmstadt, Germany, within the framework of FAIR Phase-0 (SBio-08_Quintens). L.V.E. received a PhD fellowship from the Research Fund Flanders (FWO; 11B2323N and 11B2325N) and from SCK CEN. This work was supported by a grant from the Foundation against Cancer (F/2022/2026 to F.D.S and R.Q).

## Competing interests

The authors declare no competing interests.

## References

1. Louis, D.N., et al., The 2021 WHO Classification of Tumors of the Central Nervous System: a summary. Neuro Oncol, 2021. 23(8): p. 1231-1251.10.1093/neuonc/noab106

2. Stupp, R., et al., Radiotherapy plus concomitant and adjuvant temozolomide for glioblastoma. N Engl J Med, 2005. 352(10): p. 987–96.10.1056/NEJMoa043330

3. 3. Ludwig, K., et al., *Chapter 1 - Overview of glioblastoma biological hallmarks and molecular pathology*, in Glioblastoma Resistance to Chemotherapy: Molecular Mechanisms and Innovative Reversal Strategies, R. Paulmurugan and T.F. Massoud, Editors. 2021, Academic Press. p. 1-15.

4. Wang, Q., et al., Tumor Evolution of Glioma-Intrinsic Gene Expression Subtypes Associates with Immunological Changes in the Microenvironment. Cancer Cell, 2017. 32(1): p. 42–56 e6 10.1016/j.ccell.2017.06.003

5. Verhaak, R.G., et al., Integrated genomic analysis identifies clinically relevant subtypes of glioblastoma characterized by abnormalities in PDGFRA, IDH1, EGFR, and NF1. Cancer Cell, 2010. 17(1): p. 98-110. 10.1016/j.ccr.2009.12.020

6. Patel, A.P., et al., Single-cell RNA-seq highlights intratumoral heterogeneity in primary glioblastoma. Science, 2014. 344(6190): p. 1396-401.10.1126/science.1254257

7. Richards, L.M., et al., Gradient of Developmental and Injury Response transcriptional states defines functional vulnerabilities underpinning glioblastoma heterogeneity. Nature Cancer, 2021. 2(2): p. 157–173.10.1038/s43018-020-00154-9

8. Garofano, L., et al., Pathway-based classification of glioblastoma uncovers a mitochondrial subtype with therapeutic vulnerabilities. Nature Cancer, 2021. 2(2): p. 141–156.10.1038/s43018-020-00159-4

9. Castellan, M., et al., Single-cell analyses reveal YAP/TAZ as regulators of stemness and cell plasticity in glioblastoma. Nature Cancer, 2021. 2(2): p. 174–188.10.1038/s43018-020-00150-z

10. Neftel, C., et al., An Integrative Model of Cellular States, Plasticity, and Genetics for Glioblastoma. Cell, 2019. 178(4): p. 835–849 e21. 10.1016/j.cell.2019.06.024

11. Nomura, M., et al., The multilayered transcriptional architecture of glioblastoma ecosystems. Nature Genetics, 2025. 57(5): p. 1155–1167.10.1038/s41588-025-02167-5

12. Rahman, R., et al., DNA damage response in brain tumors: A Society for Neuro-Oncology consensus review on mechanisms and translational efforts in neuro-oncology. Neuro Oncol, 2024. 26(8): p. 1367–1387. 10.1093/neuonc/noae072

13. Bao, S., et al., Glioma stem cells promote radioresistance by preferential activation of the DNA damage response. Nature, 2006. 444(7120): p. 756-60. 10.1038/nature05236

14. Tamura, K., et al., Accumulation of CD133-positive glioma cells after high-dose irradiation by Gamma Knife surgery plus external beam radiation: Clinical article. Journal of Neurosurgery JNS, 2010. 113(2): p. 310–318.10.3171/2010.2.JNS091607

15. Balbous, A., et al., A radiosensitizing effect of RAD51 inhibition in glioblastoma stem-like cells. BMC Cancer, 2016. 16: p. 604. 10.1186/s12885-016-2647-9

16. King, H.O., et al., RAD51 Is a Selective DNA Repair Target to Radiosensitize Glioma Stem Cells. Stem Cell Reports, 2017. 8(1): p. 125–139.10.1016/j.stemcr.2016.12.005

17. Kim, Y., et al., Wnt activation is implicated in glioblastoma radioresistance. Laboratory Investigation, 2012. 92(3): p. 466–473.10.1038/labinvest.2011.161

18. Wang, J., et al., Notch Promotes Radioresistance of Glioma Stem Cells Stem Cells, 2009. 28(1): p. 17–28.10.1002/stem.261

19. Tejero, R., et al., Gene signatures of quiescent glioblastoma cells reveal mesenchymal shift and interactions with niche microenvironment. EBioMedicine, 2019. 42: p. 252–269.10.1016/j.ebiom.2019.03.064

20. Osswald, M., et al., Brain tumour cells interconnect to a functional and resistant network. Nature, 2015. 528(7580): p. 93-98.10.1038/nature16071

21. Debbi, K., et al., New approaches to overcome radioresistance in glioblastoma: mechanisms, targets and role of innovative therapies, new particles and non-photon radiotherapy in 2024. A systematic review. Rep Pract Oncol Radiother, 2025. 30(2): p. 269–281.10.5603/rpor.105654

22. Carro, M.S., et al., The transcriptional network for mesenchymal transformation of brain tumours. Nature, 2010. 463(7279): p. 318-25.10.1038/nature08712

23. Yabo, Y.A., S.P. Niclou, and A. Golebiewska, Cancer cell heterogeneity and plasticity: A paradigm shift in glioblastoma. Neuro Oncol, 2022. 24(5): p. 669–682.10.1093/neuonc/noab269

24. Hubert, C.G. and J.D. Lathia, Seeing the GBM diversity spectrum. Nat Cancer, 2021. 2(2): p. 135–137.10.1038/s43018-021-00176-x

25. Chiblak, S., et al., Carbon irradiation overcomes glioma radioresistance by eradicating stem cells and forming an antiangiogenic and immunopermissive niche. JCI Insight, 2019. 4(2).10.1172/jci.insight.123837

26. Guo, Y., et al., Carbon ion irradiation conquers the radioresistance by inducing complex DNA damage and apoptosis in U251 human glioblastomas cells. Med Oncol, 2025. 42(3): p. 64. 10.1007/s12032-025-02616-5

27. Van Eupen, L., et al., Innovative radiotherapies for the treatment of glioblastoma. Neuro-Oncology Advances, 2025. 8(1). 10.1093/noajnl/vdaf255

28. Bhaduri, A., et al., Outer Radial Glia-like Cancer Stem Cells Contribute to Heterogeneity of Glioblastoma. Cell Stem Cell, 2020. 26(1): p. 48–63.e6. 10.1016/j.stem.2019.11.015

29. Couturier, C.P., et al., Single-cell RNA-seq reveals that glioblastoma recapitulates a normal neurodevelopmental hierarchy. Nat Commun, 2020. 11(1): p. 3406. 10.1038/s41467-020-17186-5

30. Winkler, F., et al., Cancer neuroscience: The past, the present, and the road ahead. Cell, 2026. 189(8): p. 2464–2489.10.1016/j.cell.2026.03.018

31. Chai, J.Y., et al., Defining the Role of GLI/Hedgehog Signaling in Chemoresistance: Implications in Therapeutic Approaches. Cancers (Basel), 2021. 13(19).10.3390/cancers13194746

32. Rajakulendran, N., et al., Wnt and Notch signaling govern self-renewal and differentiation in a subset of human glioblastoma stem cells. Genes Dev, 2019. 33(9-10): p. 498–510.10.1101/gad.321968.118

33. McCord, M. and P. Jamshidi, Targeting the cell cycle to enhance chemotherapy efficacy in glioblastoma. Neuro Oncol, 2024. 26(6): p. 1097–1098.10.1093/neuonc/noae062

34. Morrison, L., S. Loibl, and N.C. Turner, The CDK4/6 inhibitor revolution — a game-changing era for breast cancer treatment. Nature Reviews Clinical Oncology, 2024. 21(2): p. 89–105.10.1038/s41571-023-00840-4

35. Alkaç, B.Ç., et al., Therapeutic targeting of glioblastoma: miRNA signatures modulated by abemaciclib. Gene Reports, 2026. 42: p. 102413.10.1016/j.genrep.2025.102413

36. Zhao, W., et al., The CDK inhibitor AT7519 inhibits human glioblastoma cell growth by inducing apoptosis, pyroptosis and cell cycle arrest. Cell Death & Disease, 2023. 14(1): p. 11. 10.1038/s41419-022-05528-8

37. Lubanska, D. and L. Porter, Revisiting CDK Inhibitors for Treatment of Glioblastoma Multiforme. Drugs R D, 2017. 17(2): p. 255–263.10.1007/s40268-017-0180-1

38. Buckner, J.C., et al., *Radiation plus Procarbazine, CCNU,* and Vincristine in Low-Grade Glioma. N Engl J Med, 2016. 374(14): p. 1344–55.10.1056/NEJMoa1500925

39. Oehler, C., et al., Patupilone (epothilone B) for recurrent glioblastoma: clinical outcome and translational analysis of a single-institution phase I/II trial. Oncology, 2012. 83(1): p. 1–9. 10.1159/000339152

40. Stupp, R., et al., Sagopilone (ZK-EPO, ZK 219477) for recurrent glioblastoma. A phase II multicenter trial by the European Organisation for Research and Treatment of Cancer (EORTC) Brain Tumor Group. Ann Oncol, 2011. 22(9): p. 2144–2149.10.1093/annonc/mdq729

41. Dubey, J., N. Ratnakaran, and S.P. Koushika, Neurodegeneration and microtubule dynamics: death by a thousand cuts. Front Cell Neurosci, 2015. 9: p. 343. 10.3389/fncel.2015.00343

42. Iegiani, G., F. Di Cunto, and G. Pallavicini, Inhibiting microcephaly genes as alternative to microtubule targeting agents to treat brain tumors. Cell Death & Disease, 2021. 12(11): p. 956. 10.1038/s41419-021-04259-6

43. Zhou, X., et al., The Yin and Yang of Autosomal Recessive Primary Microcephaly Genes: Insights from Neurogenesis and Carcinogenesis. Int J Mol Sci, 2020. 21(5).10.3390/ijms21051691

44. Zou, Y.F., et al., Screening and authentication of molecular markers in malignant glioblastoma based on gene expression profiles. Oncol Lett, 2019. 18(5): p. 4593–4604.10.3892/ol.2019.10804

45. Ribeiro, J.H., et al., A human-specific, concerted repression of microcephaly genes contributes to radiation-induced growth defects in cortical organoids. iScience, 2025. 28(2).10.1016/j.isci.2025.111853

46. Fischer, M., Conservation and divergence of the p53 gene regulatory network between mice and humans. Oncogene, 2019. 38(21): p. 4095–4109.10.1038/s41388-019-0706-9

47. Tabnak, P., et al., Forkhead box transcription factors (FOXOs and FOXM1) in glioma: from molecular mechanisms to therapeutics. Cancer Cell International, 2023. 23.10.1186/s12935-023-03090-7

48. Maachani, U.B., et al., FOXM1 and STAT3 interaction confers radioresistance in glioblastoma cells. Oncotarget, 2016. 7(47): p. 77365–77377.10.18632/oncotarget.12670

49. Zhang, N., et al., FoxM1 inhibition sensitizes resistant glioblastoma cells to temozolomide by downregulating the expression of DNA-repair gene Rad51. Clin Cancer Res, 2012. 18(21): p. 5961–71.10.1158/1078-0432.Ccr-12-0039

50. Wang, Z., et al., Glioblastoma multiforme formation and EMT: role of FoxM1 transcription factor. Curr Pharm Des, 2015. 21(10): p. 1268–71.10.2174/1381612821666141211115949

51. Biltekin, E., et al., Inhibition of FOXM1 Leads to Suppression of Cell Proliferation, Migration, and Invasion Through AXL/eEF2 Kinase Signaling and Induces Apoptosis and Ferroptosis in GBM Cells. Int J Mol Sci, 2025. 26(14). 10.3390/ijms26146792

52. Gong, A. and S. Huang, FoxM1 and Wnt/β-catenin signaling in glioma stem cells. Cancer Res, 2012. 72(22): p. 5658–62.10.1158/0008-5472.Can-12-0953

53. Joshi, K., et al., MELK-dependent FOXM1 phosphorylation is essential for proliferation of glioma stem cells. Stem Cells, 2013. 31(6): p. 1051–63.10.1002/stem.1358

54. Zeng, W.J., et al., Aberrant ASPM expression mediated by transcriptional regulation of FoxM1 promotes the progression of gliomas. J Cell Mol Med, 2020. 24(17): p. 9613–9626.10.1111/jcmm.15435

55. Ewels, P.A., et al., The nf-core framework for community-curated bioinformatics pipelines. Nature Biotechnology, 2020. 38(3): p. 276–278.10.1038/s41587-020-0439-x

56. Love, M.I., W. Huber, and S. Anders, Moderated estimation of fold change and dispersion for RNA-seq data with DESeq2. Genome Biology, 2014. 15(12): p. 550 10.1186/s13059-014-0550-8

57. Piron, A., et al., RedRibbon: A new rank-rank hypergeometric overlap for gene and transcript expression signatures. Life Sci Alliance, 2024. 7(2).10.26508/lsa.202302203

58. Korotkevich, G., et al., Fast gene set enrichment analysis. bioRxiv, 2021: p. 060012.10.1101/060012

59. Yu, G., et al., clusterProfiler: an R package for comparing biological themes among gene clusters. Omics, 2012. 16(5): p. 284–7.10.1089/omi.2011.0118

60. Kuleshov, M.V., et al., Enrichr: a comprehensive gene set enrichment analysis web server 201C update. Nucleic Acids Research, 2016. 44(W1): p. W90–W97.10.1093/nar/gkw377

61. Garcia-Alonso, L., et al., Benchmark and integration of resources for the estimation of human transcription factor activities. Genome Research, 2019. 29(8): p. 1363–1375.10.1101/gr.240663.118

62. Alvarez, M.J., et al., Functional characterization of somatic mutations in cancer using network-based inference of protein activity. Nat Genet, 2016. 48(8): p. 838–47.10.1038/ng.3593

63. Osorio, D., et al., scTenifoldKnk: An efficient virtual knockout tool for gene function predictions via single-cell gene regulatory network perturbation. Patterns, 2022. 3(3).10.1016/j.patter.2022.100434

64. Zheng, G.X.Y., et al., Massively parallel digital transcriptional profiling of single cells. Nature Communications, 2017. 8(1): p. 14049.10.1038/ncomms14049

65. Hao, Y., et al., Dictionary learning for integrative, multimodal and scalable single-cell analysis. Nature Biotechnology, 2024. 42(2): p. 293–304.10.1038/s41587-023-01767-y

66. Chang, K., et al., The Cancer Genome Atlas Pan-Cancer analysis project. Nature Genetics, 2013. 45(10): p. 1113–1120.10.1038/ng.2764

67. Zhao, Z., et al., Chinese Glioma Genome Atlas (CGGA): A Comprehensive Resource with Functional Genomic Data from Chinese Glioma Patients. Genomics Proteomics Bioinformatics, 2021. 19(1): p. 1–12.10.1016/j.gpb.2020.10.005

68. Bowman, R.L., et al., GlioVis data portal for visualization and analysis of brain tumor expression datasets. Neuro Oncol, 2017. 19(1): p. 139–141.10.1093/neuonc/now247

69. Tang, Z., et al., GEPIA: a web server for cancer and normal gene expression profiling and interactive analyses. Nucleic Acids Res, 2017. 45(W1): p. W98–w102.10.1093/nar/gkx247

70. Cardoso-Moreira, M., et al., Gene expression across mammalian organ development. Nature, 2019. 571(7766): p. 505-509.10.1038/s41586-019-1338-5

71. Oki, S., et al., *ChIP*-*Atlas: a data*-*mining suite powered by full integration of public ChIP*-*seq data*. The EMBO Reports, 2018. 19(12): p. EMBR201846255.10.15252/embr.201846255

72. Pichol-Thievend, C., et al., VC-resist glioblastoma cell state: vessel co-option as a key driver of chemoradiation resistance. Nature Communications, 2024. 15(1): p. 3602. 10.1038/s41467-024-47985-z

73. Lippert, L., et al., PARP7 inhibition and a STING agonist potentiate radiation-induced immunogenicity in glioblastoma. Oncoimmunology, 2026. 15(1): p. 2656053. 10.1080/2162402x.2026.2656053

74. Minata, M., et al., Phenotypic Plasticity of Invasive Edge Glioma Stem-like Cells in Response to Ionizing Radiation. Cell Rep, 2019. 26(7): p. 1893–1905 e7.10.1016/j.celrep.2019.01.076

75. Al-Koussa, H., et al., The Role of Rho GTPases in Motility and Invasion of Glioblastoma Cells. Anal Cell Pathol (Amst), 2020. 2020: p. 9274016. 10.1155/2020/9274016

76. Bennan, A.B.A., et al., Joint Optimization of Photon & Carbon Ion Treatments for Glioblastoma. International Journal of Radiation Oncology, Biology, Physics, 2021. 111(2): p. 559–572.10.1016/j.ijrobp.2021.05.126

77. Chiblak, S., et al., Radiosensitivity of Patient-Derived Glioma Stem Cell 3-Dimensional Cultures to Photon, Proton, and Carbon Irradiation. Int J Radiat Oncol Biol Phys, 2016. 95(1): p. 112–119.10.1016/j.ijrobp.2015.06.015

78. Malouff, T.D., et al., Carbon Ion Therapy: A Modern Review of an Emerging Technology. Frontiers in Oncology, 2020. 10(82).10.3389/fonc.2020.00082

79. Wang, Y., et al., Charged particle therapy for high-grade gliomas in adults: a systematic review. Radiat Oncol, 2023. 18(1): p. 29.10.1186/s13014-022-02187-z

80. Abdelfattah, N., et al., Single-cell analysis of human glioma and immune cells identifies S100A4 as an immunotherapy target. Nature Communications, 2022. 13(1): p. 767.10.1038/s41467-022-28372-y

81. Hsu, J. and J. Sage, Novel functions for the transcription factor E2F4 in development and disease. Cell Cycle, 2016. 15(23): p. 3183–3190.10.1080/15384101.2016.1234551

82. Sadasivam, S. and J.A. DeCaprio, The DREAM complex: master coordinator of cell cycle-dependent gene expression. Nat Rev Cancer, 2013. 13(8): p. 585–95.10.1038/nrc3556

83. Down, C.F., et al., Binding of FoxM1 to G2/M gene promoters is dependent upon B-Myb. Biochimica et Biophysica Acta (BBA) - Gene Regulatory Mechanisms, 2012. 1819(8): p. 855–862.10.1016/j.bbagrm.2012.03.008

84. Kim, S.-H., et al., EZH2 Protects Glioma Stem Cells from Radiation-Induced Cell Death in a MELK/FOXM1-Dependent Manner. Stem Cell Reports, 2015. 4(2): p. 226–238.10.1016/j.stemcr.2014.12.006

85. Tian, J.H., et al., FOXM1-Dependent Transcriptional Regulation of EZH2 Induces Proliferation and Progression in Prostate Cancer. Anticancer Agents Med Chem, 2021. 21(14): p. 1835–1841.10.2174/1871520620666200731161810

86. Wang, Y., et al., Dysregulation of miR-C8C8-5p/FOXM1 circuit contributes to colorectal cancer angiogenesis. Journal of Experimental & Clinical Cancer Research, 2018. 37(1): p. 292. 10.1186/s13046-018-0970-5

87. Paskeh, M.D.A., et al., EZH2 as a new therapeutic target in brain tumors: Molecular landscape, therapeutic targeting and future prospects. Biomed Pharmacother, 2022. 146: p. 112532. 10.1016/j.biopha.2021.112532

88. Fan, W., H. Ma, and B. Jin, Expression of FOXM1 and PLK1 predicts prognosis of patients with hepatocellular carcinoma. Oncol Lett, 2022. 23(5): p. 146. 10.3892/ol.2022.13266

89. Fu, Z., et al., Plk1-dependent phosphorylation of FoxM1 regulates a transcriptional programme required for mitotic progression. Nat Cell Biol, 2008. 10(9): p. 1076–82. 10.1038/ncb1767

90. Gormally, M.V., et al., Suppression of the FOXM1 transcriptional programme via novel small molecule inhibition. Nature Communications, 2014. 5(1): p. 5165. 10.1038/ncomms6165

91. Ulhaka, K., et al., The Anticancer Effects of FDI-6, a FOXM1 Inhibitor, on Triple Negative Breast Cancer. Int J Mol Sci, 2021. 22(13). 10.3390/ijms22136685

92. Tian, M., et al., FOXM1 promotes the progression of non-small cell lung cancer by inhibiting miR-509-5p expression via binding to the miR-509-5p promoter region. Heliyon, 2024. 10(5): p. e27147. 10.1016/j.heliyon.2024.e27147

93. Lee, Y., et al., Characterizing and Targeting Genes Regulated by Transcription Factor MYBL2 in Lung Adenocarcinoma Cells. Cancers (Basel), 2022. 14(20). 10.3390/cancers14204979

94. Halliday, J., et al., In vivo radiation response of proneural glioma characterized by protective p53 transcriptional program and proneural-mesenchymal shift. Proc Natl Acad Sci U S A, 2014. 111(14): p. 5248–53.10.1073/pnas.1321014111

95. Fedele, M., et al., Proneural-Mesenchymal Transition: Phenotypic Plasticity to Acquire Multitherapy Resistance in Glioblastoma. International Journal of Molecular Sciences, 2019. 20(11): p. 2746

96. Bhat, K.P.L., et al., Mesenchymal differentiation mediated by NF-kappaB promotes radiation resistance in glioblastoma. Cancer Cell, 2013. 24(3): p. 331–46.10.1016/j.ccr.2013.08.001

97. Venkataramani, V., et al., Glioblastoma hijacks neuronal mechanisms for brain invasion. Cell, 2022. 185(16): p. 2899–2917.e31.10.1016/j.cell.2022.06.054

98. Bourmeau, G., et al., Proneural-mesenchymal hybrid glioblastoma cells are resistant to therapy and dependent on nuclear import. Neuro Oncol, 2025. 27(11): p. 2876–2893.10.1093/neuonc/noaf160

99. Verreault, M., et al., Preclinical Efficacy of the MDM2 Inhibitor RG7112 in MDM2-Amplified and TP53 Wild-type Glioblastomas. Clin Cancer Res, 2016. 22(5): p. 1185–96.10.1158/1078-0432.Ccr-15-1015

100. Vlashi, E., et al., Metabolic state of glioma stem cells and nontumorigenic cells. Proc Natl Acad Sci U S A, 2011. 108(38): p. 16062–7. 10.1073/pnas.1106704108

101. Shimura, T., et al., ATM-mediated mitochondrial damage response triggered by nuclear DNA damage in normal human lung fibroblasts. Cell Cycle, 2017. 16(24): p. 2345–2354. 10.1080/15384101.2017.1387697

102. Zhu, Z., et al., Zika Virus Targets Glioblastoma Stem Cells through a SOX2-Integrin α(v)β(5) Axis. Cell Stem Cell, 2020. 26(2): p. 187–204.e10. 10.1016/j.stem.2019.11.016

103. Chen, X., et al., ASPM promotes glioblastoma growth by regulating G1 restriction point progression and Wnt-β-catenin signaling. Aging (Albany NY), 2020. 12(1): p. 224–241.10.18632/aging.102612

104. Kato, T.A., et al., ASPM influences DNA double-strand break repair and represents a potential target for radiotherapy. Int J Radiat Biol, 2011. 87(12): p. 1189–95.10.3109/09553002.2011.624152

105. Li, C., et al., Identification of key modules and hub genes in glioblastoma multiforme based on co-expression network analysis. FEBS Open Bio, 2021. 11(3): p. 833–850.10.1002/2211-5463.13078

106. Liang, M.L., et al., Downregulation of miR-137 and miR-6500-3p promotes cell proliferation in pediatric high-grade gliomas. Oncotarget, 2016. 7(15): p. 19723–37.10.18632/oncotarget.7736

107. Huang, W., et al., Inhibition of KIF14 Suppresses Tumor Cell Growth and Promotes Apoptosis in Human Glioblastoma. Cell Physiol Biochem, 2015. 37(5): p. 1659–70.10.1159/000438532

108. Mathur, R., et al., Glioblastoma evolution and heterogeneity from a 3D whole-tumor perspective. Cell, 2024. 187(2): p. 446–463.e16. 10.1016/j.cell.2023.12.013

109. Gascoigne, K.E. and S.S. Taylor, Cancer Cells Display Profound Intra- and Interline Variation following Prolonged Exposure to Antimitotic Drugs. Cancer Cell, 2008. 14(2): p. 111–122.10.1016/j.ccr.2008.07.002

110. Prosser, S.L. and L. Pelletier, Mitotic spindle assembly in animal cells: a fine balancing act. Nature Reviews Molecular Cell Biology, 2017. 18(3): p. 187–201.10.1038/nrm.2016.162

111. Fischer, M., et al., Integration of TP53, DREAM, MMB-FOXM1 and RB-E2F target gene analyses identifies cell cycle gene regulatory networks. Nucleic Acids Research, 2016. 44(13): p. 6070-6086. 10.1093/nar/gkw523

112. Lee, Y., et al., FoxM1 Promotes Stemness and Radio-Resistance of Glioblastoma by Regulating the Master Stem Cell Regulator Sox2. PLOS ONE, 2015. 10(10): p. e0137703. 10.1371/journal.pone.0137703

113. Qin, Q., et al., FoxM1 knockdown enhanced radiosensitivity of esophageal cancer by inducing apoptosis. J Cancer, 2023. 14(3): p. 454–463.10.7150/jca.76671

114. Zhuang, J.-j., et al., Current strategies and progress for targeting the “undruggable” transcription factors. Acta Pharmacologica Sinica, 2022. 43(10): p. 2474–2481.10.1038/s41401-021-00852-9

115. Cai, J., et al., PROTAC: a revolutionary technology propelling small molecule drugs into the next golden age. Front Oncol, 2025. 15: p. 1676414. 10.3389/fonc.2025.1676414

116. Wang, K., et al., Peptide-based PROTAC degrader of FOXM1 suppresses cancer and decreases GLUT1 and PD-L1 expression. J Exp Clin Cancer Res, 2022. 41(1): p. 289.10.1186/s13046-022-02483-2

117. Zeng, H., et al., Self-Assembled Peptide PROTAC Prodrugs Targeting FOXM1 for Cancer Therapy. Mol Pharm, 2025. 22(6): p. 3286–3296. 10.1021/acs.molpharmaceut.5c00219

118. Madhi, H., et al., FOXM1 Inhibition Enhances the Therapeutic Outcome of Lung Cancer Immunotherapy by Modulating PD-L1 Expression and Cell Proliferation. Adv Sci (Weinh), 2022. 9(29): p. e2202702. 10.1002/advs.202202702

119. Yamashita, N., et al., FDI-6, a FOXM1 inhibitor, activates the aryl hydrocarbon receptor and suppresses tumorsphere formation. Biochem Biophys Res Commun, 2023. 639: p. 29–35. 10.1016/j.bbrc.2022.11.069

120. Bushweller, J.H., Targeting transcription factors in cancer — from undruggable to reality. Nature Reviews Cancer, 2019. 19(11): p. 611–624. 10.1038/s41568-019-0196-7

121. Swati, K., et al., Computational exploration of FOXM1 inhibitors for glioblastoma: an integrated virtual screening and molecular dynamics simulation study. Journal of Biomolecular Structure and Dynamics, 2025. 43(10): p. 5199–5217.10.1080/07391102.2024.2308772

